# *TACSTD2* expression marks the early transition to colon adenomas

**DOI:** 10.1101/2024.10.29.620817

**Authors:** Anna Siskova, Kerstin Huebner, Giulio Ferrero, Annemarie Gehring, Chuanpit Hampel, Alessio Naccarati, Barbara Pardini, Sonia Tarallo, Carol I. Geppert, Jan Prochazka, Jolana Tureckova, Jiri Jungwirth, Tomas Hucl, Jan Kral, Ludmila Vodickova, Katharina Erlenbach-Wuensch, Arndt Hartmann, Katerina Honkova, Radoslav Matej, Pavel Vodicka, Veronika Vymetalkova, Regine Schneider-Stock

**Author notes:** Correspondence to: Regine Schneider-Stock. both authors contributed equally (Anna Siskova, Kerstin Huebner). Adjunct Professor, Faculty of Pharmacy, Mahidol University, Bangkok, Thailand (Regine Schneider-Stock). both are senior authors (Veronika Vymetalkova, Regine Schneider-Stock).

## Abstract

This study aimed to address new molecular events occurring in precancerous stages of colorectal cancer (CRC) by integrated analysis of gene expression data and DNA methylation profiles.

Whole-transcriptome sequencing analysis was performed on 16 fresh frozen colorectal adenoma and matched mucosa specimens along with validation of candidates in an independent cohort of 20 fresh frozen paired adenoma and adjacent mucosa tissues as well as eight independent public datasets (335 cases). Genome-wide methylation profiles were determined for 5 adenoma pairs and confirmed by pyrosequencing on 20 tissue pairs used for validation. Functional analysis was performed *in vitro* and *in vivo* using the inflammation-associated azoxymethane/dextran sodium sulfate (AOM/DSS) and the sporadic colorectal carcinogenesis (six AOM injections) mouse models as well as *Apc^Min/+^* mice. Candidates were investigated by immunohistochemical staining of human adenomas and early-stage (pT1) CRC tumors.

A total of 1,917 differentially expressed genes and 148,191 differentially methylated CpG sites were detected in adenomas compared with adjacent mucosa samples. Based on the transcriptome data and relevance to CRC, *TACSTD2, MMP7, MMP1, CLDN2, CLDN1, and ETV4*, were selected for further validation. *TACSTD2* promoter hypomethylation in adenoma tissues was validated in 20 additional tissue pairs and corresponded with increased *TACSTD2* expression. *TACSTD2* was also overexpressed in an *in vitro* transformation model of human colonic epithelial cells. The *TACSTD2* protein TROP2 was elevated in human adenomas, pT1 tumors, and in murine adenomas, while it was absent in the unaffected adjacent mucosa. TROP2 overexpression might trigger the development of precancerous lesions and could help identify early transformation foci in colon biopsies.

**Graphical abstract:** 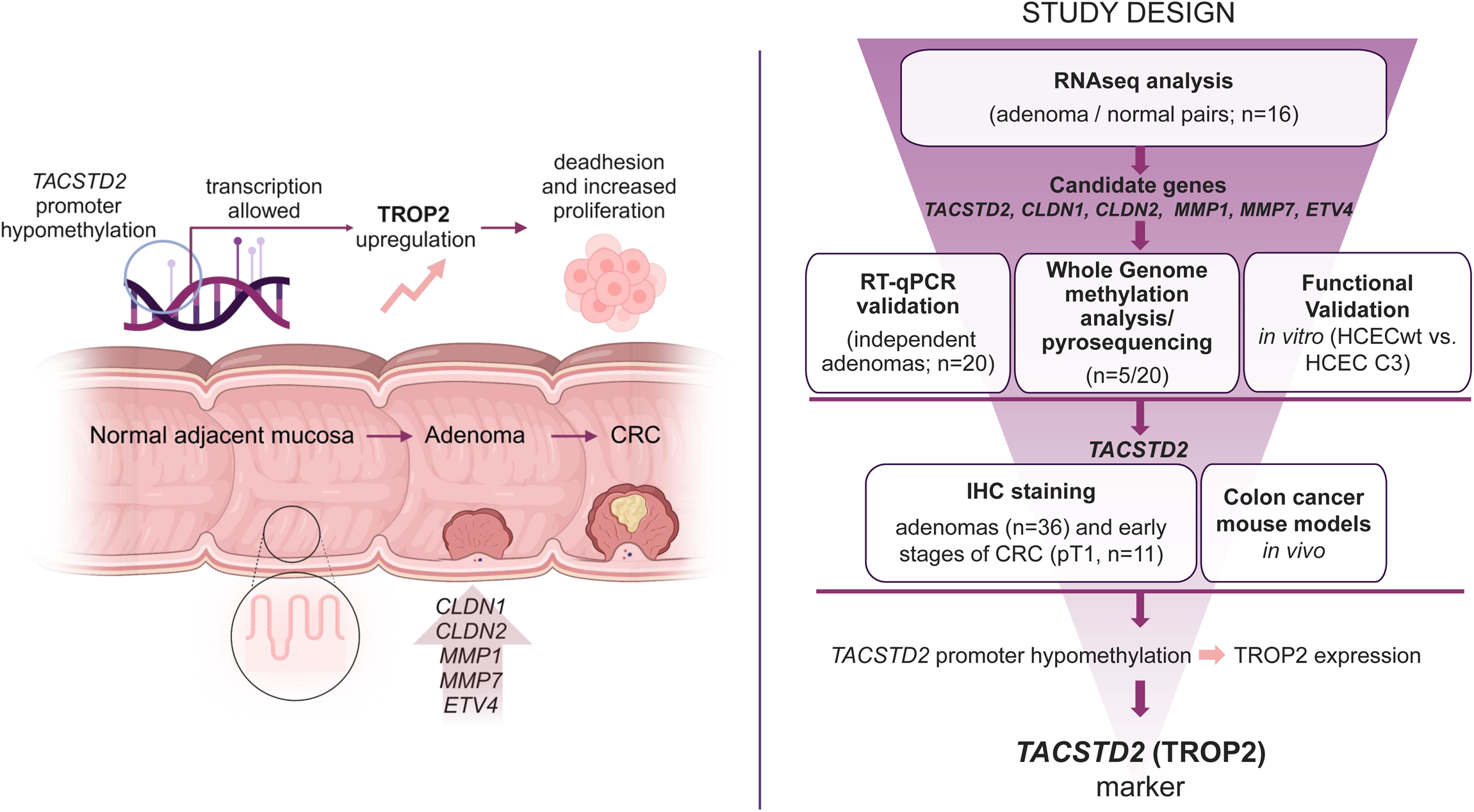

## Introduction

Colorectal adenomas are preneoplastic lesions that arise from the intestinal epithelium. If left untreated, they can progress to colorectal cancer (CRC), the second most common cause of cancer-related deaths worldwide (935,000 deaths annually) [1].

The high mortality rate of this disease is closely related to its late diagnosis at more advanced stages. Early detection of CRC or its nonmalignant precursor lesions is crucial for reducing CRC mortality [2–4]. The 5-year survival rate varies from 91-73% for patients with tumor–node– metastasis (TNM) stage I-II CRC to 13% for patients with TNM stage III-IV CRC [5, 6]. From a histological point of view, 98% of CRCs originate from adenomas [7]. Therefore, having available biomarkers associated with early CRC stages or precancerous lesions would allow early intervention with effective treatments. Several biomarkers have already been identified, including carcinoembryonic antigen (CEA) and carbohydrate antigen 19-9 (CA 19-9). These markers, although widely used for CRC detection, are not suitable for monitoring the turning points of the transition of healthy intestinal epithelium to adenoma, since they display serious limitations in terms of sensitivity and specificity [8].

The transition of adenoma into sporadic carcinoma is a gradual multistep process that may occur asymptomatically over 10-15 years or longer [9]. Most adenomatous polyps arise via the accumulation of driver mutations [9] and chromosomal instability, which promote tumor growth and the spread of cancer cells [10]. In a recent study on serrated polyps, a single-cell resolution atlas was generated indicating stem cell expansion in metaplasia [11]. Proteomic profiling revealed that metachronous advanced neoplasia could not be clearly defined by a single protein; rather, the expression pattern of a network of multiple proteins was expected to predict adenoma development [12]. To date, most studies have focused on genetic analysis of the progression of malignant colon tumors [13, 14], with less attention given to investigating cancer precursors.

In this study, RNA transcriptome analysis and methylation profiling were performed on a set of human adenoma samples to identify new drivers associated with precancerous lesion development. Several candidate genes were identified as potentially responsible for the transition from healthy tissue to adenoma, and some were further validated under *in vitro* and *in vivo* conditions. We identified *TACSTD2* (encoding for the TROP2 protein) as a potential marker with a key biological role and clinical significance in adenoma patients and potentially in early precancerous stages of CRC. We suggest that *TACSTD2* be considered a marker for adenoma development.

## Material and Methods

### Sample collection

Biological specimens and clinical and demographic data were collected from subjects recruited for hospital-based studies at the Institute for Clinical and Experimental Medicine in Prague, Czech Republic (Cohort-CZ) and Clinica S. Rita in Vercelli, Italy (Cohort-IT).

The local Ethics Committee of the Institute for Clinical and Experimental Medicine, Thomayer Hospital, Prague, Czech Republic (protocol N. G-17-03-02), and the Ethics Committee of Azienda Ospedaliera SS. Antonio e Biagio e C. Arrigo of Alessandria (Italy, protocol no. Colorectal miRNA CEC2014) approved the study. All patients provided written informed consent.

Biopsies of mucosa– adenoma pairs for Cohort-CZ (8 adenoma and 8 adjacent nontumor mucosa samples) were collected during routine screening endoscopy examinations. For Cohort-IT (8 adenoma and 8 adjacent nontumor mucosa tissues), samples were collected from all subjects during surgical resection. All samples (Cohort-CZ and Cohort-IT) were immediately transferred into cryogenic vials containing RNAlater™ Solution (Invitrogen, Carlsbad, CA) and stored at −80 °C until use. None of the patients had either a familial history of CRC or a previous personal cancer history. The clinicopathological data for all the recruited patients is reported in **Table S1**.

For further validation, a new set of samples (adenoma and adjacent mucosa pairs) was collected from an additional 20 patients recruited in the Czech Republic. Clinicopathological data for the validation set are listed in **Table S2**. The residual tissue from all adenoma samples were formalin-fixed, paraffin-embedded (FFPE). An additional set of tissue samples was collected during surgery at the Universitätsklinikum Erlangen, Germany (Cohort-GE). This set consisted of 11 FFPE tissues from early-stage (pT1) CRC patients. The Ethical Committee of the Faculty of Medicine (23-323-Br) of the Hospital Erlangen-Nürnberg approved the study. Clinical and demographic information for this cohort is reported in **Table S3**. pT1 samples were used only for immunohistochemical staining.

### RNA and DNA isolation and assessment of quality and quantity

Total RNA from Cohort-IT tissues was isolated using QIAzol (QIAGEN, Hilden, Germany) after tissue homogenization with an ULTRA-TURRAX® homogenizer (IKA-Werke GmbH & Co. KG, Staufen, Germany) followed by phenol/chloroform extraction according to the manufacturer’s standard protocol [15].

Total RNA from Cohort-CZ tissues was isolated using a RNeasy Kit (QIAGEN) employing a one-column protocol according to the manufacturer’s guidelines. Tissues were prehomogenized in a MagNa Lyser Instrument, Version 4 (Roche, Mannheim, Germany), using MagNa Lyser Green Beads (Roche). The DNase treatment procedure is described in **Supplementary Materials and Methods**.

The quality of the input RNA was determined by RNA integrity number (RIN) measurements obtained with an Agilent Bioanalyzer® RNA 6000 Nano Chip (Agilent, Santa Clara, CA). The RIN for all the samples was greater than 7. The RNA concentration was measured with a Qubit fluorometer 4.0 with the Qubit™ RNA HS and BR Assay Kits (Invitrogen).

DNA from tissues was isolated using a DNA Mini Kit (QIAGEN) following the standard protocol. The method for tissue homogenization was consistent with the method used for Cohort-CZ. The DNA concentration was measured with a Qubit fluorometer 4.0 with the Qubit™ dsDNA BR and HS Assay Kits (Invitrogen), and a NanoDrop-1000 spectrophotometer (Thermo Fisher Scientific, Waltham, MA) was used to determine the purity of the DNA (OD260/280, OD260/230).

### Transcriptomic analysis

For each sample, 500 ng of total RNA was used as the starting material for RNA-seq library preparation. RNA-seq libraries were prepared with the NEBNext® Ultra II Directional RNA Library Prep for Illumina® kits (New England Biolabs, Ipswich, MA) after ribosomal RNA depletion following the manufacturer’s instructions. The generated barcoded libraries containing fragments of approximately 300 bp (paired-end) were subjected to sequencing on the Illumina® NovaSeq™ 6000 platform (Illumina, San Diego, CA). An average of 52.2 million paired-end reads were generated for each sample (**Table S4**). A detailed description of the library preparation procedure is provided in **Supplementary Materials and Methods**.

Sequencing reads were aligned to the human GENCODE v36 transcriptome using Salmon v1.4.0 with the default settings [16]. The aligned reads were preprocessed with the *tximport* v1.18.0 [17] R package, and significantly differentially expressed genes (DEGs) were identified using *DESeq2* v1.30.0 [18] and paired Wald test [17] with adjustment for the sample cohort and adenoma subtype. Analysis was also performed by stratifying the patients based on the adenoma subtype (tubular, traditional serrated, or sessile serrated). A gene was considered a DEG if it had a median abundance value of greater than 1 transcript per million (TPM) in adenoma or adjacent mucosa samples, a Benjamini‒Hochberg (BH) adjusted *P* value of less than 0.05, and an absolute log2-fold change (FC) of greater than 1.

Functional enrichment analysis was performed using the Metascape [15], and the lists of upregulated and downregulated DEGs were used separately as input. Only the terms with a BH-adjusted *P* value <0.05 were considered significant. Data for Cohort-CZ are available in the Gene Expression Omnibus (GEO) under accession number GSE232110, while data for Cohort-IT are available upon request. Comparison of the obtained data with publicly available gene expression profiles of adenoma or CRC tissues matched with adjacent mucosa tissues was performed using GEO2R [19] with the default settings. *In silico* validation was performed with data from the following public datasets: GSE4183, GSE8671, GSE20916, GSE37364, GSE41657, GSE74602, GSE110223, and GSE110224.

### Reverse transcription and real-time quantitative PCR (RT‒qPCR)

Reverse transcription and RT‒qPCR were performed on tissue samples from the validation cohort (n=20) using a High-Capacity cDNA Reverse Transcription Kit (Thermo Fisher Scientific) starting with 800 ng of RNA. Two microliters of the 10 ng/µl cDNA suspension were used for amplification with primers specific for the target genes (*MMP7, MMP1, CLDN2, CLDN1, ETV4,* and *TACSTD2)* and the reference gene Beta2-Microglobulin (*β2-MB*) (**Table S5;** Metabion, Planegg, Germany) using the QuantiTect SYBR^®^ Green PCR Kit (QIAGEN) according to the manufacturer’s protocol. The thermocycler conditions were as follows: 5 min at 95 °C, followed by 35 cycles of denaturation for 10 sec at 95 °C and annealing for 30 sec at 60 °C. RT‒qPCR was performed on a CFX96^TM^ Real-Time System (Bio-Rad, Hercules, CA) with four technical replicates. The expression levels of target genes were normalized to those of human *β2-MB*.

FCs in gene expression were calculated as follows: FC = 2^−ΔΔCT^, where ΔΔCT is ΔCT_adenoma_ - ΔCT_mucosa_ and ΔCT is CT_target gene_ - CT_reference gene_.

### Whole genome DNA methylation

DNA (1000 ng) extracted from tissue pairs collected from five patients in Cohort-CZ was treated with sodium bisulfite using the EZ DNA Methylation Kit (Zymo Research, Irvine, CA) for the conversion of unmethylated cytosines to uracils, leaving methylated cytosines unconverted. The bisulfite-converted DNA samples were stored at −20 °C until use. Analysis of DNA methylation was performed using the Infinium Methylation EPIC v.2.0 Kit (Illumina) according to the manufacturer’s protocol and as described in Honkova et al. [20]. The use of the microarray, scanned by the iScan System (Illumina), allowed the detection of more than 850,000 methylation sites per sample across the genome at single-nucleotide resolution. The methylation status of each CpG site was estimated by comparing the intensities of the pair of methylated and unmethylated probes.

Preprocessing analyses were performed to study the distribution of beta values and the variation in methylation across all samples. Methylation data are available in GEO under accession code GSE247839. A detailed description of the methylation data processing procedure is provided in **Supplementary Methods and Materials**.

### Validation of methylated site by pyrosequencing

Details of this analysis have been described recently [21]. Briefly, for bisulfite conversion, 500 ng of DNA from the validation cohort of 20 patients (20 adenoma and 20 nontumor tissue samples) was processed with the EZ DNA Methylation-Gold Kit following the standard protocol (Zymo Research). Bisulfite conversion was followed by amplification of the target CpGs. Detailed information on the primer sequences and amplification conditions for the individual sequences of the *TACSTD2* promoter can be found in **Table S6** and **Supplementary Materials and Methods**, respectively [21].

The PCR products were prepared for pyrosequencing using the Pyromark vacuum prep tool (QIAGEN). With the corresponding sequencing primers for the four primer pairs and QIAGEN Q24 reagents, the samples were analyzed with the PyroMark Q24 system (QIAGEN). The results were evaluated using PyroMark Q24 software (QIAGEN).

### Cultivation of HCECs

Nonmalignant wild-type (wt) immortalized human colonic epithelial cells (HCECwt) (originally a gift from Dr. Pablo Steinberg (Max-Rubner Institute, Karlsruhe, Germany, Herbst et al. 2006 [22])) and their derived cell line HCEC C3 were cultured in HCEC medium (PAN Biotech, Aidenbach, Germany) supplemented with 2 mM L-Glutamine (GlutaMax, Gibco Life Technologies, Grand Island, NY), 30 µg/ml bovine pituitary extract (PromoCell, Germany), 38 µg/ml ascorbic acid (Sigma–Aldrich, Darmstadt, Germany), 1 nM dexamethasone (Sigma– Aldrich), and 100 nM retinoic acid (Sigma–Aldrich).

Since HCEC medium generally does not contain fetal bovine serum for trypsin neutralization, cells were cultured on Corning® CellBIND® surface culture dishes (Merck, Darmstadt, Germany), centrifuged after trypsinization, and resuspended in fresh medium to avoid the effects of residual trypsin (PAN Biotech). The HCEC cell line was genotyped using the Multiplex Cell Authentication service by Multiplexion (Heidelberg, Germany) and was regularly tested and confirmed to be negative for mycoplasma contamination.

### Generation of pre-transformed epithelial cells via anchorage blockade

Seven thousand and five hundred HCEC cells were seeded in HCEC medium in standard 10 cm cell culture dishes coated with 1% agarose and incubated at 37 °C with 5% CO_2_ for 96 h. Then, the cells were collected and seeded in one well of a 12-well cell culture plate without an agarose coating. Surviving cells (hereafter referred to as HCEC C1 cells) adherent to the plate were retained, and the nonadherent cells were washed away and discarded. The cells were further cultured in cell culture dishes without an agarose coating. When the number of cells was sufficient, the anchorage blockade cycle was repeated two more times (3 cycles in total) to generate HCEC C3 cells. The steps for Immunofluorescence assays of proliferation and RNA isolation with clean-up, reverse transcription, and RT‒qPCR are described in detail in **Supplementary Materials and Methods**.

### Mouse models

All mice were bred and housed in the Czech Centre for Phenogenomics (Prague, Czech Republic), and the *in vivo* procedures were performed under protocols approved by the Animal Care Committee of the Institute of Molecular Genetics (Prague, Czech Republic, license number AVCR 1653-2022 SOV II).

### Mouse models of sporadic colorectal carcinogenesis

To mimic the sporadic colorectal carcinogenesis process, we chose two available models in the present work—induction after long-term exposure of C57BL/6NCrl mice to AOM and the *Apc*^min/+^ mouse model.

For the long-term tumorigenesis assay, eight-week-old C57BL/6NCrl mice were injected intraperitoneally with AOM (10 mg/kg) at weekly intervals for a total of six injections. The animals were then sacrificed 8 months later (week 38), and the intestinal tubes were dissected and processed for further histological analysis of lesions.

For the *Apc*^min/+^ model, 15-23-week-old *Apc*^min/+^ mice (of mixed sex), with mutations in the *Apc* gene were sacrificed, and both the small and large intestines were harvested and fixed for histological analysis.

### Azoxymethane/dextran sulfate sodium (AOM-DSS) mouse model of colorectal carcinogenesis

To study AOM-DSS-induced tumorigenesis, eight- to twelve-week-old male C57BL/6NCrl mice were injected intraperitoneally with AOM (10 mg/kg; Sigma–Aldrich) in PBS. Seven days later, the first of two cycles of *ad libitum* exposure to 2% DSS (TdB Consultancy, Sweden) in the drinking water was started. Each cycle lasted for five days and was separated by a 14-day recovery period. The second cycle of DSS exposure was followed by a four-week period of tumorigenesis. Six animals were kept on untreated water as controls.

After the mice were sacrificed (at the experimental endpoint, 64 days or earlier due to animal welfare ethics), the colon tubes were longitudinally cut, washed thoroughly with PBS, rolled, fixed with 4% formaldehyde for 24 h, transferred to 70% alcohol, and finally embedded in paraffin. Immunohistochemical staining of mouse tissues is described in **Supplementary Materials and Methods**.

## Results

### Whole-transcriptome sequencing revealed significantly different expression profiles between adenomas and the adjacent mucosa

From the differential expression analysis (discovery set of samples, n=16 patients), a total of 1,917 DEGs (adjusted *P <*0.05, |log2FC| > 1, **Fig. 1a**) were identified between adenoma and the paired adjacent mucosa samples. Among these DEGs, 688 were upregulated and 1,229 were downregulated in adenoma samples (**Table S7a**). The analysis was also performed in relation to the histological type of adenoma (**Table S1**) to determine any possible difference among adenoma subtypes compared with their paired adjacent mucosa: 1,876 DEGs were found for tubular adenoma, 1,455 DEGs were found for traditional serrated adenoma, and 237 DEGs were found for sessile serrated adenoma.

**Figure 1.**
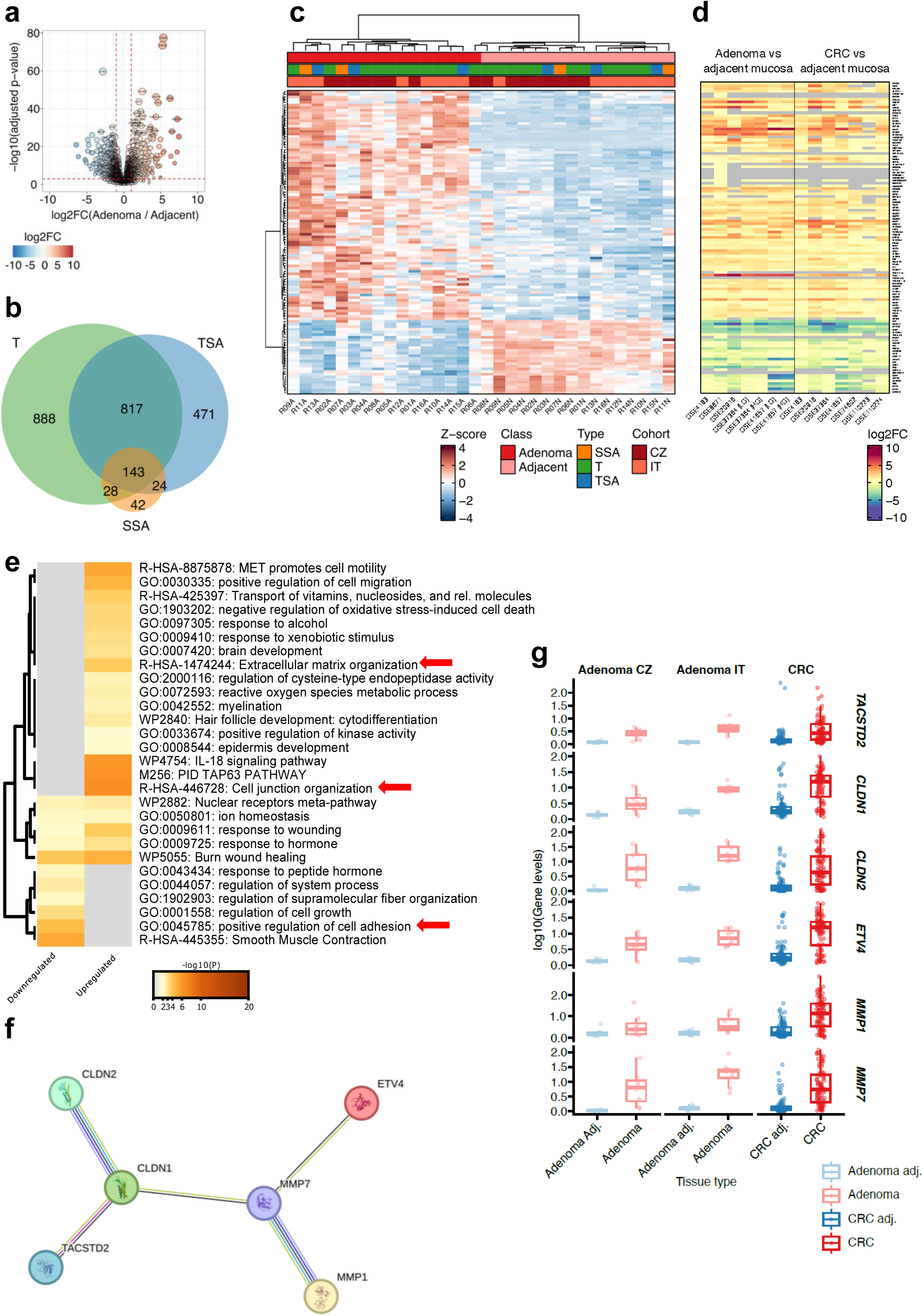
Six candidate genes were selected based on transcriptomic analysis of colorectal adenomas. **a**. Volcano plot showing the log2FC and adjusted *P* values resulting from differential expression (DE) analysis between adenoma and adjacent colonic mucosa in the discovery set of samples (n=16 patients). Thresholds: adjusted *P* value *<*0.05 and |log2FC| > 1. The dot size is proportional to the -log10 (adjusted *P* value). **b.** Venn diagram showing the overlap between differentially expressed genes (DEGs) identified in the analysis of discovery set of samples stratified by adenoma type (T=tubular adenoma; TSA=traditional serrated; SSA=sessile serrated). **c.** Heatmap showing the Z scores of the normalized DEGs identified as commonly dysregulated in different adenoma types with respect to the paired adjacent mucosa samples (T=tubular adenoma; TSA=traditional serrated; SSA=sessile serrated). **d.** Heatmap of the log2FC values computed *in silico* between the levels of the DEGs in adenoma or CRC tissues compared to those in the matched colonic mucosa. For two datasets (GSE37364 and GSE41657), the analysis was performed separately for LGAs and HGAs. **e.** Top functional terms enriched in the 115 DEGs. The red arrows indicate dysregulated pathways related to the chosen candidate genes. **f.** STRING database analysis of interactions among the six candidate genes selected for validation. **g.** Expression levels of the candidate genes selected for the adenoma signature (*MMP7, MMP1, CLDN1, CLDN2, ETV4, TACSTD2*) as measured in the adenoma–adjacent tissue pairs from Cohort-CZ and Cohort-IT and in colon carcinomas from Gagliardi et al. 2023 in comparison with the surrounding adjacent mucosa.

Since the aim of the study was to search for markers that would be consistent and valid regardless of the histological subtype of adenoma, the study focused on the DEGs common to all three adenoma subtypes (143 DEGs) **(Fig. 1b)**. Among these DEGs, 115 still exhibited significant differential expression and most of these (n=87, 75.6%) were upregulated in adenoma tissues (**Fig. 1c**). The most significantly upregulated DEG in adenoma tissue was *MMP7* (log2FC = 7.46, combined *adj. P = 5.19e-26*). On the other hand, the most significantly downregulated DEG was *MS4A12* (log2FC = −3.38, *adj. P = 1.63e-6*). The normalized expression levels of the 115 DEGs are reported in **Fig. 1c** and **Table S7b.** The results were also validated *in silico* in a panel of independent public datasets reporting expression profiles for adenomas and adjacent unaffected mucosa as well as for carcinomas and adjacent unaffected mucosa (**Fig. 1d** and **Table S7c**). Again, the results of these analyses confirmed the observed differential expression in both adenoma and CRC samples compared with the paired adjacent mucosa samples (**Fig. 1c-d**).

Functional enrichment analysis of the 115 DEGs (**Table S8a**) was performed to identify the related biological functions (**Table S8b**). The most significant (*adj. P <*0.05) functional terms are reported in **Fig. 1e**. Specifically, GO terms related to cell adhesion (GO:0045785), matrix organization (R-HSA-1474244) and specific signaling pathways, including the c-MET (R-HSA-8875878), Interleukin-18 (IL-18) (WP4754), and Tumor protein 63 (TAp63) signaling pathways (M256) were enriched in the upregulated DEGs. Conversely, terms related to cell adhesion and regulation of cell growth, including regulation of system process (GO:0044057) and regulation of supramolecular fiber organization (GO:1902903), were enriched in the downregulated DEGs.

### Selection of candidate genes and validation of these genes based on high-throughput data analysis

We focused on genes displaying increased expression (upregulation) in adenoma compared to adjacent mucosa samples. The selection of the total number of genes to be validated was constrained by the quantity of RNA obtained from adenoma biopsies, which was a limiting factor. Consequently, validation was performed on six genes (*MMP7*, *MMP1*, *CLDN2*, *CLDN1*, *ETV4*, and *TACSTD2*) chosen from the 20 most upregulated genes (**Table 1**) and based on enrichment analysis and current research literature on CRC.

**Table 1.**
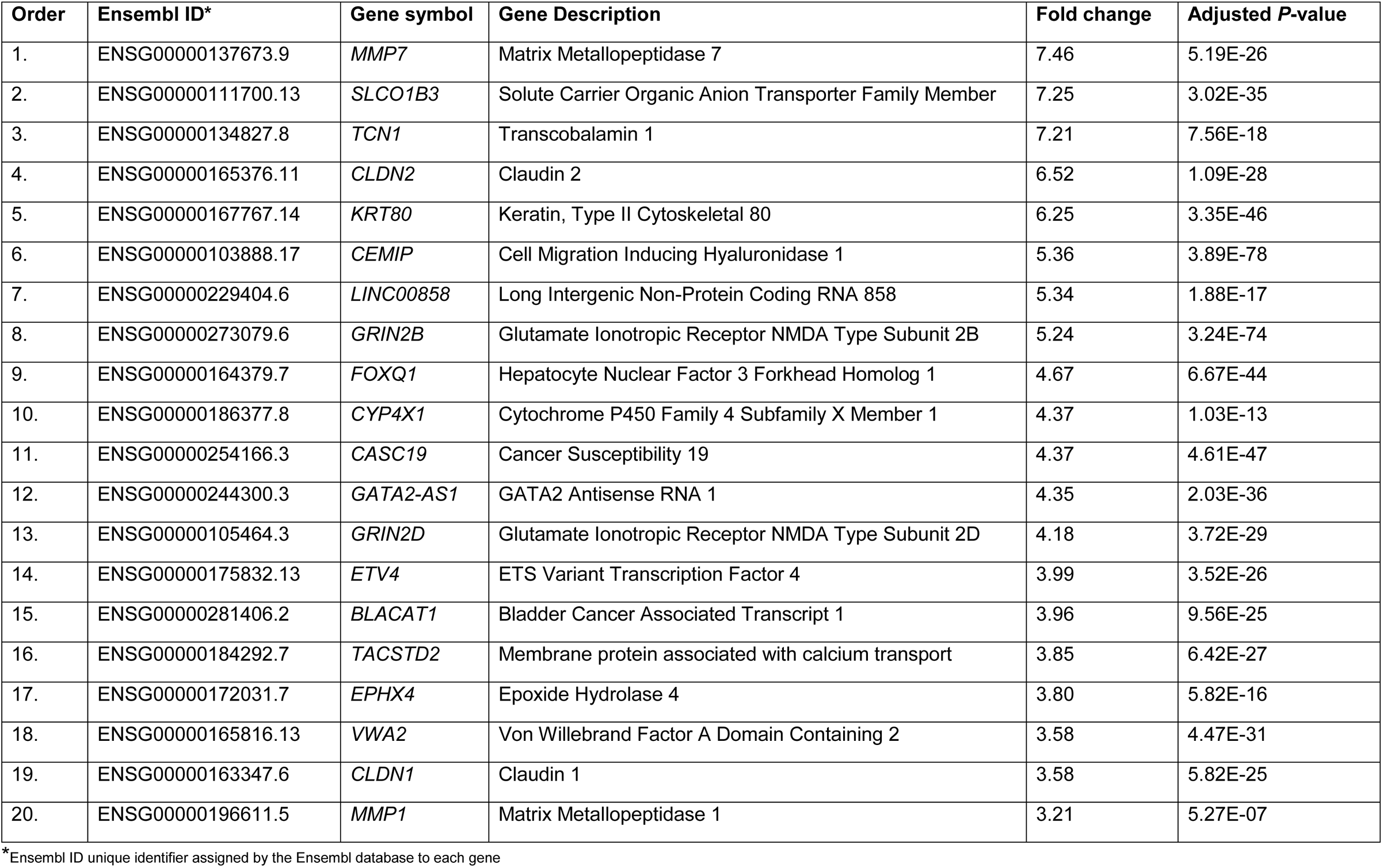
Top 20 most up-regulated genes from RNA-seq analysis of 16 patients.

| Order | Ensembl ID* | Gene symbol | Gene Description | Fold change | Adjusted *P*-value |
| --- | --- | --- | --- | --- | --- |
| 1. | ENSG00000137673.9 | *MMP7* | Matrix Metallopeptidase 7 | 7.46 | 5.19E-26 |
| 2. | ENSG00000111700.13 | *SLCO1B3* | Solute Carrier Organic Anion Transporter Family Member | 7.25 | 3.02E-35 |
| 3. | ENSG00000134827.8 | *TCN1* | Transcobalamin 1 | 7.21 | 7.56E-18 |
| 4. | ENSG00000165376.11 | *CLDN2* | Claudin 2 | 6.52 | 1.09E-28 |
| 5. | ENSG00000167767.14 | *KRT80* | Keratin, Type II Cytoskeletal 80 | 6.25 | 3.35E-46 |
| 6. | ENSG00000103888.17 | *CEMIP* | Cell Migration Inducing Hyaluronidase 1 | 5.36 | 3.89E-78 |
| 7. | ENSG00000229404.6 | *LINC00858* | Long Intergenic Non-Protein Coding RNA 858 | 5.34 | 1.88E-17 |
| 8. | ENSG00000273079.6 | *GRIN2B* | Glutamate Ionotropic Receptor NMDA Type Subunit 2B | 5.24 | 3.24E-74 |
| 9. | ENSG00000164379.7 | *FOXQ1* | Hepatocyte Nuclear Factor 3 Forkhead Homolog 1 | 4.67 | 6.67E-44 |
| 10. | ENSG00000186377.8 | *CYP4X1* | Cytochrome P450 Family 4 Subfamily X Member 1 | 4.37 | 1.03E-13 |
| 11. | ENSG00000254166.3 | *CASC19* | Cancer Susceptibility 19 | 4.37 | 4.61E-47 |
| 12. | ENSG00000244300.3 | *GATA2-AS1* | GATA2 Antisense RNA 1 | 4.35 | 2.03E-36 |
| 13. | ENSG00000105464.3 | *GRIN2D* | Glutamate Ionotropic Receptor NMDA Type Subunit 2D | 4.18 | 3.72E-29 |
| 14. | ENSG00000175832.13 | *ETV4* | ETS Variant Transcription Factor 4 | 3.99 | 3.52E-26 |
| 15. | ENSG00000281406.2 | *BLACAT1* | Bladder Cancer Associated Transcript 1 | 3.96 | 9.56E-25 |
| 16. | ENSG00000184292.7 | *TACSTD2* | Membrane protein associated with calcium transport | 3.85 | 6.42E-27 |
| 17. | ENSG00000172031.7 | *EPHX4* | Epoxide Hydrolase 4 | 3.80 | 5.82E-16 |
| 18. | ENSG00000165816.13 | *VWA2* | Von Willebrand Factor A Domain Containing 2 | 3.58 | 4.47E-31 |
| 19. | ENSG00000163347.6 | *CLDN1* | Claudin 1 | 3.58 | 5.82E-25 |
| 20. | ENSG00000196611.5 | *MMP1* | Matrix Metallopeptidase 1 | 3.21 | 5.27E-07 |

\*Ensembl ID unique identifier assigned by the Ensembl database to each gene

Among the candidate genes selected for validation, *MMP7* was chosen because it was the most highly upregulated gene in the entire transcriptome datasets. Since *MMP1* belongs to the family of matrix metalloproteinases [23], which also includes *MMP7*, and the extracellular matrix (ECM) organization pathway was one of the pathways with the greatest enrichment in the upregulated DEGs (**Fig. 1e**; red arrow), both genes were selected for validation. *CLDN2* and *CLDN1*, whose products are transmembrane proteins that enable the formation of cellular tight junctions [24, 25], also participate in one of the most highly upregulated pathways in adenoma tissue (**Fig. 1e**; red arrow). The transcription factor *ETV4* was also selected for validation based on its role in promoting the upregulation of matrix metalloproteinases [26] and its association with tumor progression in various cancer types [27]. The final candidate, *TACSTD2* (coding for a protein known as TROP2), has already been well described to play a role in the progression and metastasis of various cancers, including breast cancer and CRC [28, 29]. Interestingly, TROP2 and CLDN1 have already been shown to be direct binding partners [30]. Both might affect cell adhesion, a biological pathway that was found to be enriched in the downregulated genes in the enrichment analysis (**Fig. 1e**; red arrow). In addition to the functional properties of these six candidates and their participation in cancer processes (**Table S9**), interactions among them were also detected *in silico* (**Fig. 1f**).

Our results were also validated *in silico* in an independent public dataset [17] with expression profiles for adenoma and adjacent normal tissues as well as for carcinoma and adjacent normal tissues. The analysis confirmed the same trend of upregulated expression in both adenoma and CRC samples [15] (**Fig. 1g**). Interestingly, the gene expression levels in the mucosa adjacent to the carcinoma tissues were greater than those in the mucosa adjacent to the adenoma tissues, suggesting that the adjacent tissue was already transformed but did not exhibit visible morphological alterations (**Fig. 1g**).

RT‒qPCR analysis of the mRNA expression of *MMP7, MMP1, CLDN2, CLDN1, ETV4,* and *TACSTD2* in an independent validation cohort of 20 adenoma patients revealed that the average log2FC of all these genes was significantly greater (*P <*0.05) in adenomas than in the adjacent colonic mucosa (**Fig. S1a-b**). This observation confirmed the results obtained from the RNA-seq analysis.

### Genome-wide methylation assay reveals relevant differences between adenomas and adjacent mucosa

To better understand the regulatory mechanism underlying the increased gene expression, the CpG methylation profiles were explored at the genome-wide level in five available DNA samples from paired adenoma and adjacent mucosa tissues already included in the RNA-seq analysis.

There were significant differences between the methylation profiles of adenoma and unaffected tissues in all patients (**Fig. 2a**). According to the data analysis, 148,191 CpG sites were differentially methylated in adenomas compared to the adjacent mucosa (**Fig. S2a**). Among the differentially methylated CpG sites in adenomas, 23,178 were hypermethylated and 125,013 were hypomethylated (**Fig. S2b**). The comparison was based on the analyzed Δβ values and CpG sites with an *adj. P <*0.05.

**Figure 2.**
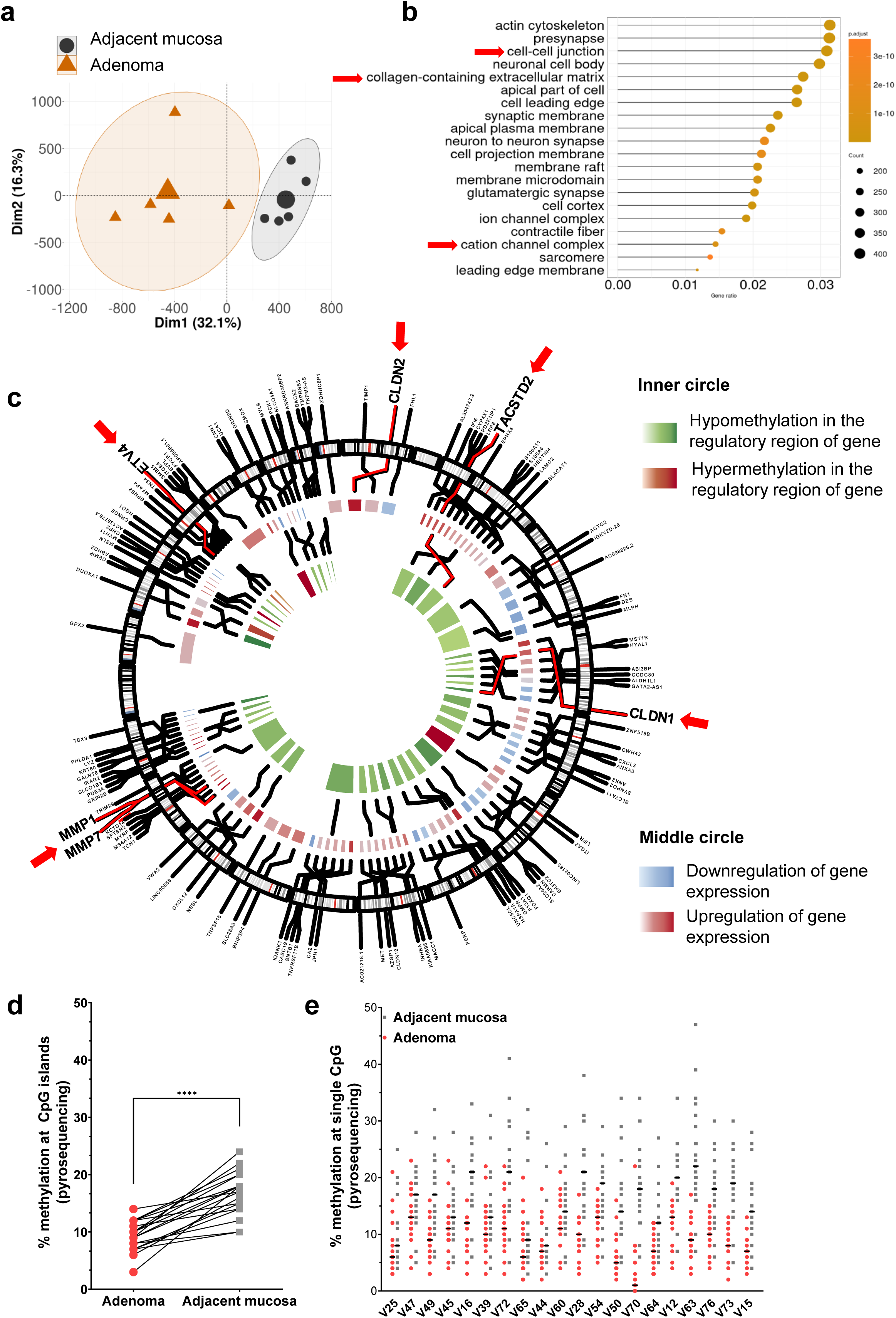
Hypomethylation of the *TACSTD2* promoter was found in adenoma tissue compared to adjacent mucosa tissues. **a.** Principal component analysis (PCA) plot based on the differential DNA methylation of CpG loci showing the separation between adenoma and adjacent tissues. **b.** List of the Gene Ontology (GO) pathways most significantly enriched in the set of differentially methylated genes. The red arrows indicate the pathways related to the chosen candidate genes. **c.** Circos plot integrating the transcriptome sequencing data with the gene promoter methylation profile data. The red arrows indicate the six candidate genes whose expression was independently validated in 20 adenoma patients. Only *CLDN1* and *TACSTD2* exhibited hypomethylation of their promoter regions (inner circle). **d.** Summary graph of pyrosequencing data showing significant (**** *P <*0.0005) hypomethylation in adenoma samples compared to adjacent colonic mucosa samples from all 20 patients in the validation cohort. **e.** Comparison of the percentage of *TACSTD2* promoter methylation in adenoma versus adjacent colonic mucosa samples (pyrosequencing, validation cohort; n=20). General hypomethylation was observed in the promoter region in the adenoma samples from each patient.

Enrichment analysis revealed that the functional pathways containing the most highly differentially methylated genes also contained the six selected candidate genes: cell-to-cell junction (*CLDN1*, *CLDN2*), ECM organization (*MMP1*, *MMP7*), and calcium signaling pathway (*TACSTD2*) (**Fig. 2b**; red arrows). Interestingly, hypomethylation of the gene promoter region (inner green circle in **Fig. 2c**) was detected only in the *TACSTD2* (**Fig. S2c** and **Fig. S2d**) and *CLDN1* (**Fig. S2e**) genes, explaining their increased expression. For the *ETV4*, *MMP1,* and *MMP7* genes (**Fig. S2f-h**), no hypomethylation in the regulatory regions was observed, and no probes for the *CLDN2* gene were present in the microarray. Since *TACSTD2* promoter hypermethylation has recently been described in colon carcinoma [21], hypomethylation in the promoter region of *TACSTD2* was further investigated (**Fig. S2c).** Pyrosequencing of the *TACSTD2* promoter region confirmed that the methylation levels in adenomas were lower than those in the corresponding adjacent mucosa in the validation cohort of 20 patients (**Fig. 2d-e).**

### *In vitro* validation of candidate genes

Since anchorage-independent growth with loss of cell attachment is one of the first features of the precancerous stage [31] and the candidate genes have a functional connection to the adhesion pathway, the levels of the six candidate genes were measured in an *in vitro* model using HCECwt cells and the corresponding pre-transformed HCEC C3 cells generated by repeated adhesion blockade cycles (**Fig. 3a**). Indeed, HCEC C3 cells were characterized by the formation of 3D clusters mimicking cluster rosettes (**Fig. 3a**). Moreover, greater proliferative activity was observed for HCEC C3 cells than for HCECwt cells (**Fig. 3b-d**). Thus, HCEC C3 cells might be a suitable “natural” model for early precancerous transformation without the need to use carcinogens or mutational drivers. There was a significant increase (*P <*0.05) in the mRNA expression levels of *TACSTD2* and *ETV4* in HCEC C3 cells compared to HCECwt cells, as measured by RT‒qPCR. *CLDN1* was downregulated in HCEC C3 cells compared with HCECwt cells (*P <*0.05), while *CLDN2* mRNA expression did not significantly differ between the two cell lines (*P >*0.05) (**Fig. 3e**). *MMP1* was not expressed in either cell line, whereas *MMP7* was expressed solely in HCECwt cells. Data are not displayed on the graphs for *MMP1* and *MMP7* genes.

**Figure 3.**
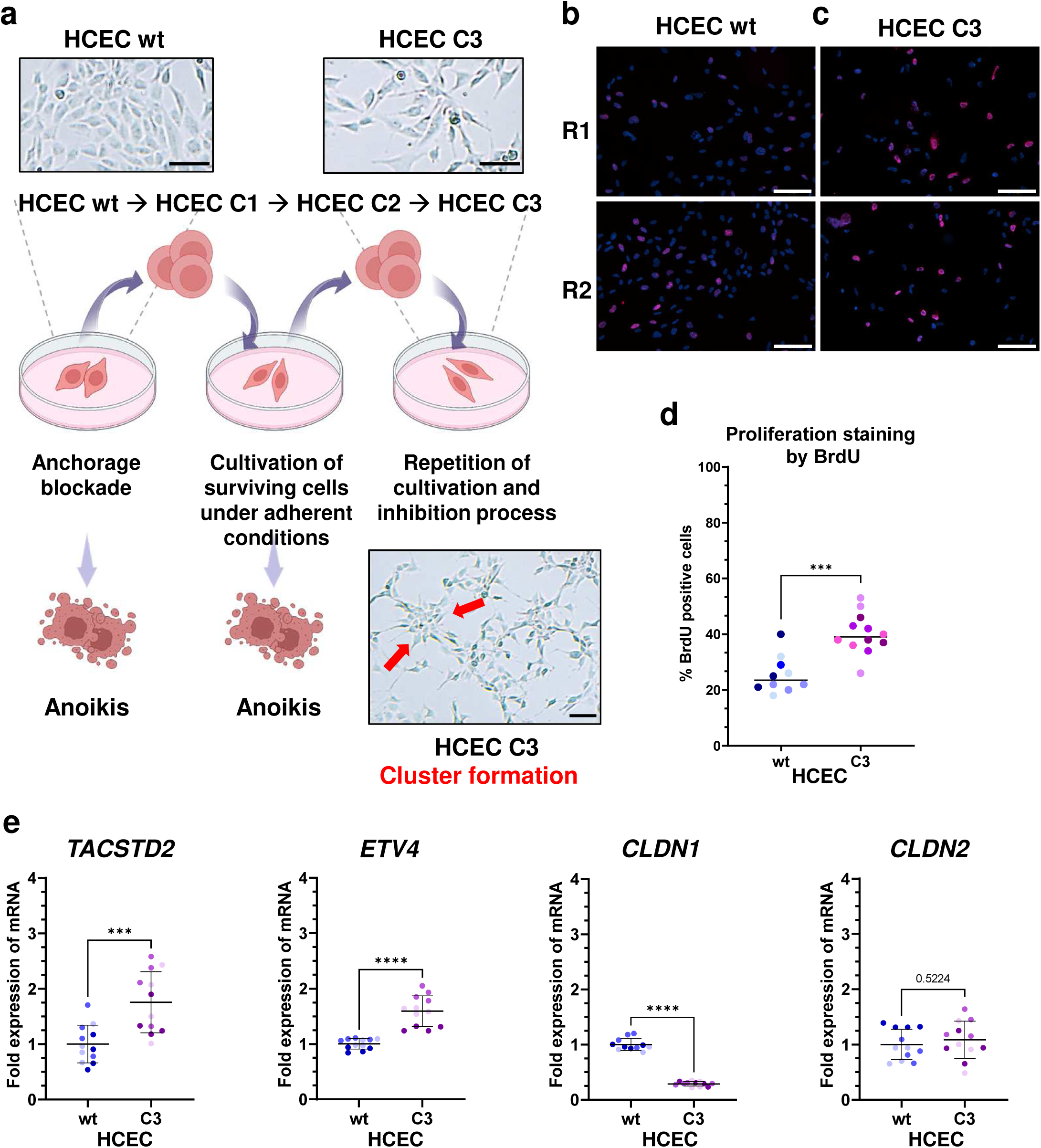
*TACSTD2* was upregulated in a cell line model mimicking colorectal adenoma cells relative to wild-type cells representing healthy intestinal mucosa. **a.** Generation of HCEC C3 cells from wild-type (wt) HCECs by repeated cycles of adhesion blockade and culture. HCEC C3 cells were characterized by cluster formation (red arrows). Scale: 100 µm. Schematic diagram was created with Biorender.com. **b.** Proliferation assay by 5-bromo-2’-deoxyuridine (BrdU) incorporation in HCECwt cells; R1= biological replicate 1; R2 = biological replicate 2. The pinkish-red color indicates a nucleus in an actively dividing cell (S phase of the cell cycle), a marker of proliferation. The blue color (4’,6-diamidino-2-phenylindole (DAPI) staining) indicates the nuclei of all cells regardless of the cell cycle phase. Scale: 100 µm **c.** Proliferation assay by BrdU incorporation in HCEC C3 cells. In both biological replicates (R1 and R2), more proliferating cells were visible than were observed in HCECwt cells (Panel **b**). Scale: 100 µm. **d.** Proliferation assay by BrdU incorporation in HCEC C3 and HCECwt cells. Proliferation was increased in HCEC C3 cells compared to HCECwt cells (*** *P <*0.005). **e.** Comparison of candidate gene expression levels in HCEC C3 and HCECwt cells. *TACSTD2* (*** *P <*0.005) and *ETV4* (**** *P <*0.0005) were upregulated in HCEC C3 cells, *CLDN1* (**** *P <*0.0005) was downregulated in HCEC C3 cells, and *CLDN2* (*P >*0.05) did not change in expression between the two cell lines. *MMP7* and *MMP1* were not expressed, data are not displayed on the graphs.

### TROP2 is a marker for the precancerous stage of CRC

Thus far, our findings strongly support the previous observations that *TACSTD2* (TROP2) is closely functionally linked to fetal progenitor cells, adhesion, and proliferation [28, 32]. Thus, we focused on TROP2 for further validation at the translational level.

Immunostaining for TROP2 and the proliferation marker Ki67 was performed on all 36 available FFPE adenomas (16 adenomas from sequencing analysis and 20 adenomas from the validation cohort) and on a new set of 11 pT1 colon tumors (Cohort-GE) (**Table S3**). Immunohistochemical staining for TROP2 was detected in all adenoma tissues, regardless of histological type (**Fig. 4a-c),** location, or frequency (**Fig. 4d)**. The adjacent mucosa was largely negative for TROP2 staining (**Fig. 4a-c**). The Ki67 and TROP2 expression indices were slightly greater in high-grade adenomas (HGAs) than in low-grade adenomas (LGAs), although the difference was not significant (**Fig. 4e)**. Moreover, the TROP2 expression index was not significantly increased in pT1 samples. However, the expression index of the proliferation marker Ki67 was slightly greater at pT1 (**Fig. 4f**).

**Figure 4.**
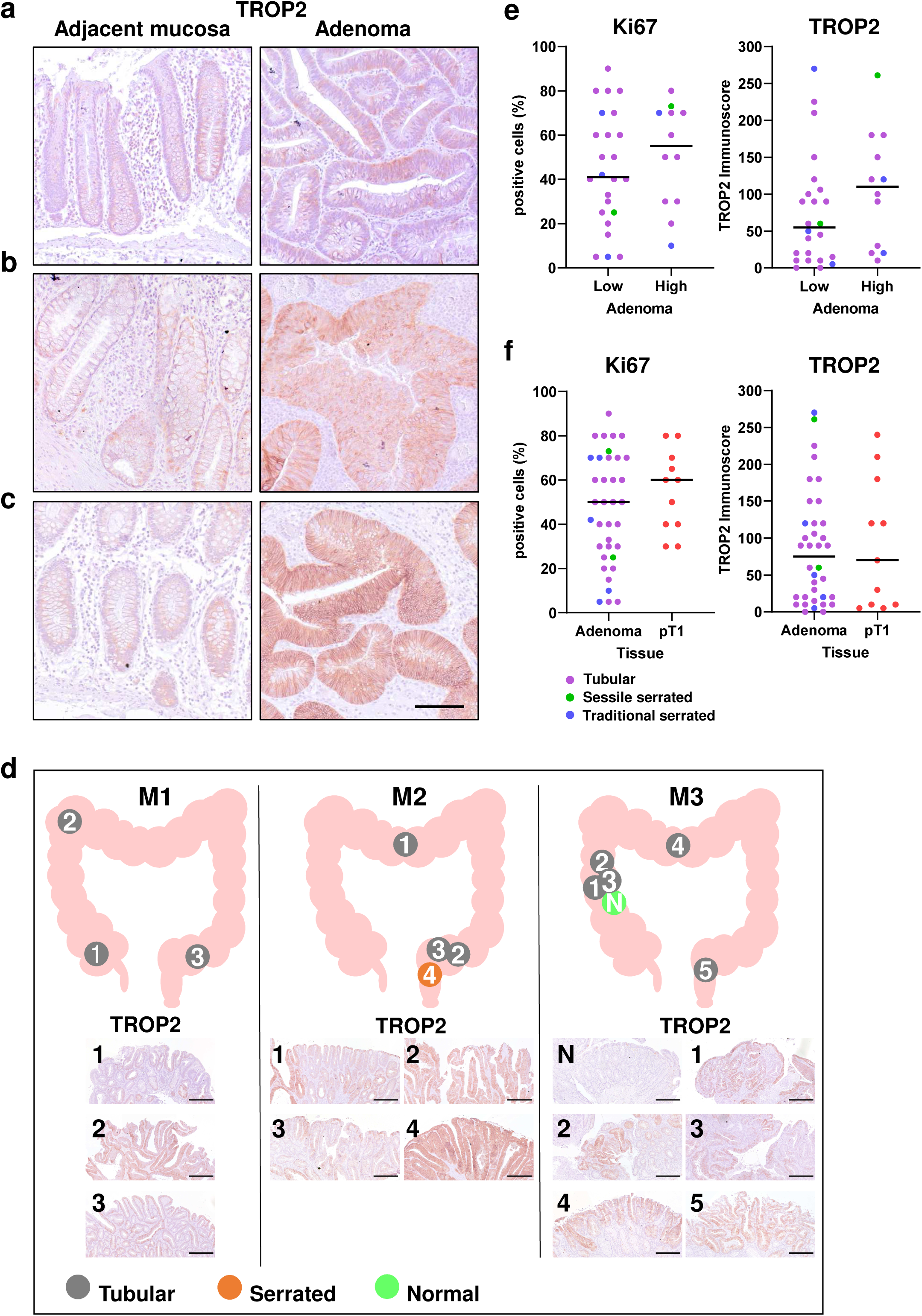
TROP2 was upregulated in human colorectal adenomas regardless of their histological type or localization. **a-c.** TROP2 immunohistochemical (IHC) staining in three representative types of adenomas ((**a**) tubular, (**b**) sessile serrated, (**c**) traditional serrated) and the paired adjacent mucosa sample from the same patient. Scale: 100 µm **d.** TROP2 expression in adenomas was found to be independent of the location, frequency, and histological type in three independent patients from Italy with multiple adenomas. Patient **M1** had three tubular adenomas: in the caecum (**1**), flexura hepatica (**2**), and sigmoid colon (**3**). Patient **M2** had three tubular adenomas and one sessile serrated adenoma: in the transverse colon (**1**), sigmoid colon (**2** and **3**), and rectum (**4**). Notably, there was increased visual expression in the sessile serrated adenoma compared to the tubular adenomas. Patient **M3** had five tubular adenomas: in the ascending colon (**1-3**), transverse colon (**4**), and rectosigmoid colon (**5**). N represents reference unaffected adjacent mucosa tissue without TROP2 expression. Scale: 300 µm. **e.** Comparison of IHC staining for Ki67 (left) and TROP2 (right) in LGAs and HGAs. **f.** Comparison of IHC staining for Ki67 (left) and TROP2 (right) in all adenomas versus pT1 tumors.

### Increased Trop2 expression *in vivo* in adenomas from different colon cancer mouse models

To further analyze the role of Trop2, the murine homolog of TROP2, in colon adenomas, three different mouse colon cancer models were evaluated: the AOM-DSS model, the sporadic 6xAOM model and the *Apc^min/+^* mouse model (**Fig. 5a-c**, **Fig. S3**). We observed mostly LGAs and HGAs (with tubular-villous histology) in the colon in all three models, with *Apc^min/+^* mice developing adenomas predominantly in the small intestine (jejunum and ileum) (**Fig. 5a, Fig. S3**). Very few carcinomas were visible in the AOM-based models. In general, partially cribriform HGA of the colonic mucosa with a dense arrangement of dysplastic cells and a papillary exophytic growth pattern without an invasive character were observed in all the models. The adenoma cells were hyperchromatic and pleomorphic with loss of polarization to the basal orientation. Notably, in line with the progression of dysplasia, the subpopulation of goblet cells was lost in HGAs, as reflected by the loss of Periodic acid–Schiff (PAS) staining, whereas the adjacent colonic mucosa was still PAS positive. As expected, an increase in the Ki67 proliferation index was observed in adenomas compared to the adjacent colonic mucosa (**Fig. 5a-c**).

**Figure 5.**
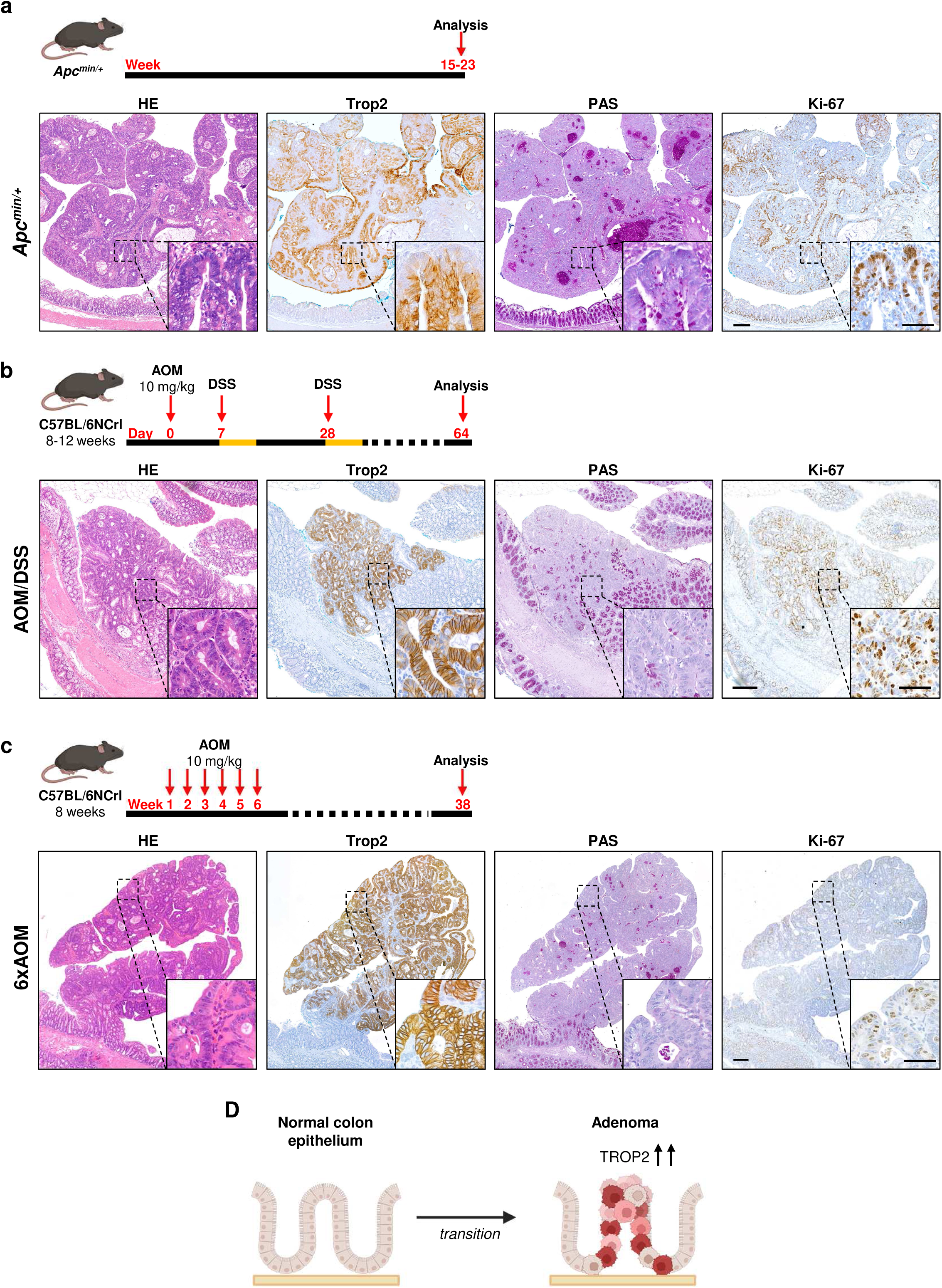
Trop2 is upregulated in colon adenomas formed in sporadic and colitis-associated mouse models. **a.** *Upper panel*: Experimental setup for the induction of sporadic colorectal carcinogenesis using *Apc^min/+^* mice. *Lower panel*: Representative images of HE-, Trop2-, PAS-, and Ki-67-stained adenomas in the colon (n=10). **b.** *Upper panel*: Experimental setup for the induction of inflammation-associated colorectal carcinogenesis (AOM-DSS model). C57BL/6NCrl mice were treated with an intraperitoneal injection of 10 mg/kg AOM and exposure to 2 % DSS at the indicated time points. *Lower panel*: Representative images of HE-, Trop2-, PAS-, and Ki-67-stained adenomas in the colon (n=3). **c.** *Upper panel*: Experimental setup for the induction of sporadic colorectal carcinogenesis (6xAOM model). C57BL/6NCrl mice were treated with a total of six intraperitoneal injections of 10 mg/kg AOM at the indicated time points. *Lower panel*: Representative images of HE-, Trop2-, PAS-, and Ki-67-stained adenomas in the colon (n=2). **a-c.** Trop2 upregulation is associated with a decrease in PAS positivity and an increase in the number of Ki-67-positive neoplastic cells. The adjacent normal mucosa retains PAS positivity, and Ki-67 staining is only present in the crypt base. Overview, scale: 200 µm; inset, scale: 50 µm. **d.** Schematic model of TROP2 upregulation during the transition of healthy colonic mucosa to dysplastic adenoma. All schematic diagrams were created with Biorender.com.

All adenomas, independent of histology, were characterized by marked upregulation of Trop2 (**Fig. 5a-c, Fig. S3**) compared to that in the unaffected adjacent normal intestinal epithelium.

## Discussion

This study presents novel findings related to the relatively underexplored topic of precancerous stages of CRC. We aimed to identify early triggers that could promote the transition of a healthy colon epithelium to adenoma, a precancerous lesion of CRC, and consequently lead to the development of CRC.

Using whole-transcriptome and methylome analyses, we identified six genes (T*ACSTD2, MMP7, MMP1, CLDN2, CLDN1,* and *ETV4)* related to the transition from healthy mucosa to adenoma. The increased expression and the methylation profiles of these genes were also validated in independent cohorts of adenoma patients.

We selected TROP2 for further translational and functional *in vitro* and *in vivo* analyses since it has frequently been reported to be highly expressed in different types of cancers originating from the epithelium, such as CRC [33–35]. Although TROP2 is recognized as a novel promising drug target in breast cancer [36, 37], its role in CRC is still highly debated. In CRC, increased TROP2 expression is associated with tumor progression and metastasis and modulates intercellular adhesion [28]. In agreement with its marked upregulation in early adenoma, TROP2 has been shown to play a role in embryonic development, maintenance of the epithelial barrier, and damage repair [37]. As a transmembrane glycoprotein, it mediates calcium signaling and seems to be indispensable for cell proliferation and survival [38]. In this context, TROP2 is known to bind directly to another candidate protein, CLDN1, thus serving as an anchor or transporter during CLDN1 rearrangement or acting as a stabilizer to prevent claudin degradation [30]. CLDN1, a main component of tight junctions, is essential for ensuring cell adhesion to the surrounding ECM. In addition, MMP1 and MMP7 are functionally dependent on the calcium level, which is implicitly affected by TROP2 [38].

Recently, Svec et al. [39] reported high TROP2 expression in *Apc^min/+^* mouse-derived adenomas. They separated cancerous epithelial cells according to the abundance of the TROP2 protein and conducted enrichment analysis specifically within the TROP2-rich cell population. The KRAS and NF-κB signaling pathways exhibited the highest scores, and “ECM organization” and “Regulation of cell proliferation” were the most significantly enriched Gene Ontology biological processes [39]. Notably, we also identified these pathways as important for adenoma development—for example, the ECM organization pathway, containing the *MMP7* and *MMP1* genes, and the cell-to-cell junction pathway, containing the *CLDN2* and *CLDN1* genes. *ETV4*, which encodes a transcription factor of the Erythroblast Transformation Specific (ETS) family, plays a role in proliferation as well as roles in differentiation and migration [40]. In our study, we discovered that TROP2 was upregulated not only in the adenomas of *Apc^min/+^*mice but also in inflammation-associated adenomas generated in the AOM-DSS model and sporadic adenomas generated in the AOM model. The carcinogen AOM is known to induce Wnt signaling by inducing β-Catenin mutations [41], and the resulting adenomas show signs of dedifferentiation due to the loss of goblet cells. Conversely, the Wnt signaling pathway increases the expression of claudins to affect selective epithelial permeability and tight junctions [42].

Usually, healthy epithelial cells undergo anoikis, a form of cell death caused by loss of adhesion blockade. Under physiological conditions, anoikis prevents dysplasia. Increased proliferation results in the outgrowth and displacement of cells in the intestinal epithelium, with loss of adequate contact between the cells and the ECM. Thus, high TROP2 expression might represent a survival mechanism activated upon loss of attachment to the ECM and provide a selection advantage to the cell subpopulation when cell contacts are affected due to increased proliferation. The observation of the robust nonadherent growth of TROP2-overexpressing colon tumors [28] suggests that TROP2 might promote the proliferation of newly transformed clones by creating space for cell division. Accordingly, TROP2 has been considered as a stem/progenitor marker and a trigger of self-renewal and hyperplasia in prostate basal cells, bladder cells, and intestinal progenitor cells [35, 37]. In the context of intestinal tissue, which is subjected to continuous renewal of luminal stem cells that differentiate into intestinal epithelial cells, the accumulation of epigenetic changes and mutations within intestinal stem cells could lead to sustained TROP2 overexpression.

*In vitro,* HCEC C3 cells, which had undergone various numbers of cycles of substrate adhesion blockade and had not been subjected to genetic manipulation or treatment with carcinogens, grew in 3D clusters independent of the substrate. These cells exhibited increased proliferation and increased *TACSTD2* expression. However, unlike *ETV4* and *TACSTD2*, the remaining candidate genes displayed opposite expression patterns in HCEC C3 cells and HCECwt cells (as observed for *CLND1*), showed no significant difference in expression between the HCECwt and the HCEC C3 cells (as observed for *CLDN2*), or were either not expressed (*MMP1*) or expressed solely in the HCECwt cells (*MMP7*). We hypothesize that the lack of expression or varied expression patterns of the other candidate genes (*MMP7, MMP1, CLDN2* and *CLDN2*) in the *in vitro* model might be due to a deficiency of the intercellular matrix for degradation under these experimental conditions. Although HCECs may serve as an acceptable “natural” model for intestinal cells, the results of repeated cycles of adhesion blockade suggest that the *in vitro* conditions might not sufficiently mirror the extracellular environment of the intestinal mucosa. However, this finding strengthens our hypothesis that TROP2 is upregulated as an intrinsic feature of epithelial cells and seems not to be heavily dependent on the ECM.

*TACSTD2* promoter demethylation has been recently identified as an epigenetic regulatory mechanism underlying the increased TROP2 expression in CRC [21]. We also detected promoter demethylation in early adenomas by whole-methylome and pyrosequencing analyses. Although every adenoma showed clear TROP2 upregulation, some advanced-stage CRCs again exhibited loss of TROP2, an observation that is not fully understood but could be explained at least partially by the high intratumor heterogeneity of late-stage tumors [28]. TROP2 was detected in adenomas regardless of the histological type, frequency, or location, whereas it was absent in the adjacent mucosa, making it an appropriate marker for precancerous and early stages of CRC.

Finally, our study provides substantial insights into the role of TROP2 in colon adenoma development. We suggest that TROP2 overexpression plays a pivotal role in the initial stages of colon cancer development. The validation of TROP2 expression in colon biopsies could help identify early transformation foci.

## Conclusions

We discovered a novel gene expression signature associated with colon adenoma development. TROP2 triggers early colon epithelial cell transformation. Thus, TROP2 expression in colon biopsies could aid in identifying early transformation foci. Whether TROP2 is also a marker for predicting progression to cancer requires further investigation.

## Statements & Declarations

### Funding

The work was supported by the Czech Science Foundation (GACR) (grant no. 22-05942S) and the Programme Exceles LX22NPO5102. The work was funded partly by a grant from the Deutsche Forschungsgemeinschaft: project no. 468812580 to RSS. AG was funded by the Interdisciplinary Center of Clinical Research (IZKF) at the Universitätsklinikum Erlangen. This project has received funding from the European Union’s Horizon 2020 Research and Innovation Program under grant agreement no. 825410 (ONCOBIOME project to Alessio Naccarati, Barbara Pardini, and Sonia Tarallo). The research leading to these results has received funding from AIRC under IG 2020 - ID 24882 (PI: Alessio Naccarati) to Alessio Naccarati.

This publication reflects only the authors’ views, and the European Commission is not responsible for any use that may be made of the information it contains.

### Autor contributions

Anna Siskova, MSc (Conceptualization: Lead; Data curation: Lead; Methodology: Lead; Project administration: Supporting; Validation: Lead; Writing – original draft: Lead, Writing – review & editing: Lead)

Kerstin Huebner, PhD (Data curation: Equal; Methodology: Equal; Validation: Equal; Writing – original draft: Supporting; Writing – review & editing: Equal)

Giulio Ferrero, PhD (Data curation: Lead; Formal analysis: Lead; Software: Lead; Visualization: Equal; Writing – original draft: supporting; Writing – review & editing: Supporting)

Annemarie Gehring, MD (Data curation: Supporting; Methodology: Supporting; Software: Supporting; Validation: Supporting)

Chuanpit Hampel, PhD (Methodology: Equal; Supervision: Supporting; Validation: Supporting)

Alessio Naccarati, PhD (Conceptualization: Supporting; Funding acquisition: Supporting; Resources: Supporting; Supervision: Supporting; Writing – review & editing: Lead)

Barbara Pardini, Ph.D. (Conceptualization: Supporting; Funding acquisition: Supporting; Resources: Supporting; Supervision: Supporting; Writing – review & editing: Lead)

Sonia Tarallo, PhD (Data curation: Supporting; Methodology: Supporting; Validation: Supporting; Writing – review & editing: Supporting)

Carol Geppert, PD MD (Data curation: Supporting; Investigation: Supporting; Validation: Supporting)

Jan Prochazka, PhD (Methodology: Supporting; Resources: Supporting; Validation: Supporting)

Jolana Tureckova PhD (Data curation: Supporting; Validation: Supporting; Writing – review & editing: Supporting)

Jiri Jungwirth, MD (Data curation: Supporting; Resources: Supporting)

Tomas Hucl, Prof., MD (Data curation: Supporting; Resources: Supporting)

Jan Kral, MD, PhD (Data curation: Supporting; Resources: Supporting)

Ludmila Vodickova, MD, CSc. (Funding acquisition: Supporting; Project administration: Supporting; Resources: Supporting)

Katharina Erlenbach-Wuensch, PD MD (Methodology: Supporting; Resources: Supporting; Validation: Supporting)

Arndt Hartmann, Prof., MD (Funding acquisition: Supporting; Project administration: Supporting; Resources: Equal)

Katerina Honkova, PhD (Formal analysis: Equal; Methodology: Equal; Validation: Supporting; Visualization: Supporting)

Radoslav Matej, Prof., MD (Data curation: Supporting; Resources: Supporting)

Pavel Vodicka, CSc., MD (Conceptualization: Supporting; Funding acquisition: Equal; Project administration: Supporting; Resources: Supporting; Supervision: Equal Writing – review & editing: Equal)

Veronika Vymetalkova, PhD (Conceptualization: Lead; Funding acquisition: Lead; Investigation: Supporting; Project administration: Equal; Resources: Supporting; Supervision: Lead; Writing – original draft: Supporting; Writing – review & editing: Lead)

Regine Schneider-Stock, Prof. (Conceptualization: Lead; Funding acquisition: Equal; Investigation: Supporting; Methodology: Supporting; Project administration: Lead; Resources: Lead; Supervision: Lead; Visualization: Supporting; Writing – original draft: Supporting; Writing – review & editing: Lead)

### Data availability

The datasets generated during and/or analyzed the current study are available in:

Methylation profiling: https://www.ncbi.nlm.nih.gov/geo/query/acc.cgi?acc=GSE247839; GSE247839

RNAseq: https://www.ncbi.nlm.nih.gov/geo/query/acc.cgi?acc=GSE232110; GSE232110

### Ethics approval

All procedures conformed to the Helsinki Declaration for research on humans. The local Ethics Committee of the Institute for Clinical and Experimental Medicine, Thomayer Hospital, Prague, Czech Republic (protocol N. G-17-03-02), and the Ethics Committee of Azienda Ospedaliera SS. Antonio e Biagio e C. Arrigo of Alessandria (Italy, protocol no. Colorectal miRNA CEC2014) approved the study. All patients provided written informed consent.

An additional set of tissue samples was collected during surgery at the Universitätsklinikum Erlangen, Germany. The Ethical Committee of the Faculty of Medicine (23-323-Br) of the Hospital Erlangen-Nürnberg approved the study.

### Consent to participate

All patients provided written informed consent.

### Consent for publication

All the authors agree to publish this paper.

## Acknowledgment

The authors thank Sarah Schmitt and Christa Winkelmann for their excellent technical help. The present work was performed in partial fulfillment of the requirements for obtaining the degree Dr. med. for AG at the Universitätsklinikum Erlangen at Friedrich-Alexander Universität Erlangen-Nürnberg.

