## Supplementary Materials and Methods for "*TACSTD2* expression marks the early transition to colon adenomas"

**Supplementary Methods**

**Supplementary Figure legends**

**Supplementary Tables**

### Supplementary Methods

#### DNase treatment

To remove all traces of DNA, before RNA-seq library preparation, total RNA from tissue samples from Cohort-IT was cleaned up and DNase-treated with the RNA Clean & Concentrator™-5 kit (Zymo Research) following manufacturer's protocol. For Cohort-CZ, DNase treatment was performed by RNase-Free DNase Set (Qiagen, Germany).

#### RNA-seq library preparation

An aliquot with 1000 ng of total RNA was used for cDNA libraries preparation according to the NEBNext Ultra II Directional RNA Library Prep Kit for Illumina (New England Biolabs, USA), as provided by the manufacturer. Ribosomal RNAs (rRNAs) were removed using NEBNext rRNA Depletion Kit (Human/Mouse/Rat, New England Biolabs, USA). After RNA heat fragmentation, hybridization with random primers, and purification, first cDNA strand was reversely transcribed, followed by second cDNA strand incorporating dUTP for directionality, thus enabling strand-specific library construction. To ensure an efficient adaptor ligation, the ends of the cDNA molecules were prepared through terminal repair and single adenylation at 3' end. Following the excision of the uracil containing strand, adapter-ligated fragments were PCR enriched with unique indexed primers NEBNext Multiplex Oligos for Illumina (Unique Dual Index Primer Pairs, New England Biolabs, USA).

The quality and the size distribution of the cDNA library was checked by Agilent High Sensitivity DNA chip (Agilent Technologies, USA), and was run on Agilent Bioanalyzer 2100 High sensitivity DNA Kit (Agilent Technologies, USA).

For the accurate quantification of the DNA library, the samples were analyzed by using the KAPA Library Quantification Kit for Illumina, a fluorometric based system (Thermo Fisher Scientific Baltics, Lithuania).

**Processing of methylation data**

Raw microarray data were downloaded as idat files, imported to the R environment, and processed with the minfi package [1]. Data were normalized using the Funnorm method. Beta values for the determination of the levels of methylation were calculated using the minfi package as the ratio of the fluorescent signals from the methylated vs. unmethylated sites. A series of filtering was performed. Probes with SNPs at CpG sites and the cross-reactive probes were also excluded to obtain the resulting number of 794,441 probes [2]. We estimated the associations between principal components and slide factors and used the Combat function (sva package) for batch correction [3].

Principal components analysis (PCA) was performed to identify the variance using the covariance matrix. We identified differentially methylated loci using the top Table function (limma package). A CpG was defined as differentially methylated with a Benjamini–Hochberg-adjusted (false discovery rate (FDR)) p-value lower than 0.05 [4]. The list of genes associated to differentially methylated CpGs/regions were analyzed with DAVID to investigate the Gene Ontology (GO) terms enriched.

The proportions of genomic regions to gene positions were analyzed using the annotatr package [5]. The annotation of the CpG site to ENTREZID, the plots of the GO pathways, and an enrichment map were obtained using clusterProfiler package v4.0 [6].

**Amplification of target CpGs in pyrosequencing method**

Two µl of BCD aliquots were amplified by PyroMark PCR Kit (Qiagen, Germany). Four different pairs of primers were used for *TACSTD2* promoter sequence: one Qiagen PyroMark CpG Assay PCR primers, and three forward and reverse self-designed primers using the Pyromark Assay Design Software (Qiagen, Germany). The products of the amplification were determined in 2.5% agarose gel with 15 µl Roti-Safe GelStain (CarlRoth, Germany), 5 µl sample, and 5 µl of 100bp DNA Ladder (Nippon Genetics Europe GmbH, Germany). Gel electrophoresis was performed for 40 minutes at 80V.

**Immunofluorescence proliferation assay in *in vitro* studies**

Twenty thousand cells for HCECwt and HCEC C3 cell lines were each seeded into 8-well cell chamber (Ibidi, Germany) with 300µl medium per well and cultured for 48 h. After reaching 70% confluency, cells were incubated at 37°C for 6 h with 0.03 mg/ml 5-Bromo-2´-Deoxyuridine (BrdU), (Merck, Germany), subsequently fixed with 70% methanol and processed as described in the Immunofluorescence Protocol for primary antibody BrdU (Bu20a) Mouse mAb (#5292, 1:500, Cell Signaling, USA). Goat anti-Mouse Alexa Fluor™ Plus 555 (#A32727, 1:500, Invitrogen, USA) was used as a secondary antibody. Nuclei were visualized by DAPI (#MBD0015, 1:1000, Sigma Aldrich, Germany).

Images were taken by Nikon Eclipse Ti-S (Nikon, Japan) at 20× magnification and percentage of BrdU positive cells was estimated in ImageJ software (National Institutes of Health, USA).

**RNA isolation with clean up, Reverse transcription, RT-qPCR in *in vitro* studies**

Total RNA from cell pellets of HCECwt and HCEC C3 cells was isolated using QIAzol® Lysis Reagent (Qiagen) combined with RNeasy Mini Kit (Qiagen, Germany) and RNase-Free DNase Set (Qiagen, Germany) according to the manufacturer's protocol.

Reverse transcription was performed with 2 µg of total RNA using the QuantiTect Reverse Transcription Kit (Qiagen, Germany) according to the manufacturer's protocol. Final cDNA was diluted to 10 ng/µl. RT-qPCR was performed according to the same procedure as it is described in main text.

**Immunohistochemical staining of mouse tissue in *in vivo* studies**

Formalin-fixed and paraffin embedded (FFPE) swiss roles were cut into 1-3 µm thin sections and subsequently deparaffinized according to standard immunohistochemical (IHC) procedures. Hematoxylin and eosin (HE) as well as PAS stainings were performed using established protocols of the routine laboratory of the Institute of Pathology in Erlangen (University Hospital Erlangen, Germany). IHC staining for TROP2 (1:2000, ab214488, Abcam, UK) and Ki67 (1:2000, #12202, Cell Signaling, USA) were manually conducted. For this, heat-induced antigen retrieval was performed in a pressure cooker at up to 120°C for 1 min in citrate, pH6 (TROP2) and TRS, pH6 (Ki67) buffers, respectively. After blocking of endogenous peroxidase activity, sections were incubated with primary antibodies overnight at room temperature (RT). Then, slides were incubated with biotinylated secondary antibodies (Vector Laboratories, USA) and detected with VECTASTAIN® Elite® ABC Kit (Vector Laboratories, USA) and 3,3'-diaminobenzidine tetrahydrochloride (DAB) substrate (Dako/ Agilent Technologies, USA). Sections were counterstained with hematoxylin. Stained swiss role sections were then scanned with a Panoramic MIDI or Panoramic 250 Flash system (Camera type: CIS VCC-FC60FR19CL; objective: Plan-Apochromat; magnification: 40x; Camera adapter magnification: ×1; 3DHISTECH, Hungary) and documented with the CaseViewer software (version 2.4, 3DHISTECH, Hungary).

### Supplementary Figure legends

**Supplementary Figure 1.** Expression levels of the six candidate genes (*MMP7, MMP1, CLDN1, CLDN2, ETV4, TACSTD2*) measured by RT-qPCR in an independent validation group of 20 patients with adenomas (18 Tubular and 2 Traditional serrated). Red bars show the fold change expression of each mRNA in adenoma tissue compared to the adjacent for each patient.

**Supplementary Figure 2. a.** Distribution and subclassification of the total DNA methylation probes detected (n=865,859) and those differentially methylated in adenomas (A) compared to the adjacent mucosa (N) (n=148,151). **b.** Proportion of significant hyper- (n=23,178) and hypo- (n=125,013) methylated CpG sites in adenoma versus adjacent mucosa according to the genome position. **c.** Heatmap of methylation profiles of *TACSTD2* gene in five adenoma patients from the Cohort-CZ (discovery group). Significant hypomethylation can be observed for adenoma samples in correspondence of cg06149158 in the regulatory region (red rectangle). Green color describes hypomethylation while red color depicts hypermethylation. **d-h.** Plots showing the differentially methylation levels (expressed as Beta-values) among the 5 adenoma and adjacent tissue pairs for all the CpG islands in *TACSTD2* (**d**), *CLDN1* (**e**), *ETV4* (**f**), *MMP1* (**g**), and *MMP7* (**h**) genes. There were no probes for *CLDN2* gene to generate the plot.

**Supplementary Figure 3.** Trop2 is upregulated in adenomas of ileum and jejunum in a mouse models for sporadic colorectal carcinogenesis. *Upper panel*: Experimental setup for the development of sporadic colorectal carcinogenesis using $Apc^{min}$ mice. *Middle panel*: Representative images of HE-, Trop2-, PAS-, and Ki-67-stained adenomas in the ileum (n=9). *Lower panel*: Representative images of HE-, Trop2-, PAS-, and Ki-67-stained adenomas in the jejunum (n=9). In both intestinal sections, Trop2 upregulation is associated by a loss of PAS positivity and an increase in Ki-67-positive neoplastic tumor cells. The adjacent normal mucosa retains PAS positivity and Ki-67 staining is only present in the crypt base. Overview, scale: 200 µm; insert, scale: 50 µm.

### Supplementary Tables

**Supplementary Table S1. Patient characteristics (n=16) and association with immunohistochemical expression of TROP2 and Ki67 in adenomas.**

| ID sample | Cohort | Gender | Age (years) | Histology type of adenoma* | TROP2 Score** | Ki67 Score | Grade | Localization |
|---|---|---|---|---|---|---|---|---|
| R01 | Czech | Female | 66 | T | 60% / 3 | 50% | High | colon |
| R02 | Czech | Female | 61 | T | 70% / 3 | 70% | Low | colon |
| R03 | Czech | Male | 60 | TSA | 10% / 2 | 10% | High | colon |
| R04 | Czech | Female | 56 | T | 60% / 2 | 20% | Low | rectum |
| R05 | Czech | Female | 43 | T | Negative | 60% | Low | rectum |
| R06 | Czech | Male | 64 | T | 15% / 3 | 80% | Low | colon |
| R07 | Czech | Male | 63 | SSA | 20% / 3 | 25% | Low | colon |
| R08 | Czech | Male | 67 | T | 5% / 3 | 15% | Low | colon |
| R09 | Italy | Male | 56 | T | 30% / 3 | 20% | High | colon |
| R10 | Italy | Male | 70 | T | 10% / 1 | 25% | Low | colon |
| R11 | Italy | Female | 64 | SSA | 87% / 3 | 73% | High | rectum |
| R12 | Italy | Male | 62 | T | 53% / 2 | 33% | Low | colon |
| R13 | Italy | Female | 74 | TSA | 90% / 3 | 42% | Low | colon |
| R14 | Italy | Male | 63 | T | 10% / 2 | 5% | Low | colon |
| R15 | Italy | Female | 69 | TSA | 50% / 1 | 5% | Low | colon |
| R16 | Italy | Male | 69 | T | 90% / 1 | 5% | Low | colon |

*T = Tubular, TSA = Traditional serrated, SSA = Sessile serrated; **% of positive tumor cells / staining intensity (0-3)

**Supplementary Table S2. Patient characteristics (Validation Cohort; n=20) and association with immunohistochemical expression of TROP2 and Ki67 in adenomas.**

| ID sample | Gender | Age (years) | Histology type of adenoma* | TROP2 Score** | Ki67 Score | Grade | Localization |
|---|---|---|---|---|---|---|---|
| V28 | Female | 69 | T | 10% / 2 | 80% | Low | colon |
| V39 | Male | 52 | T | 50% / 2 | 30% | Low | rectum |
| V44 | Male | 44 | T | N*** / 0 | 40% | Low | colon |
| V45 | Male | 58 | T | 30% / 3 | 50% | Low | colon |
| V47 | Male | 63 | T | 5% / 2 | 90% | Low | colon |
| V49 | Male | 61 | TSA | 5% / 1 | 70% | Low | colon |
| V50 | Male | 52 | T | 30% / 1 | 80% | High | colon |
| V54 | Male | 60 | T | 50% / 2 | 70% | High | rectum |
| V60 | Female | 70 | T | 20% / 1 | 70% | High | colon |
| V63 | Female | 50 | T | 5% / 2 | 60% | High | colon |
| V64 | Female | 42 | T | 60% / 3 | 50% | High | colon |
| V65 | Male | 56 | T | 50% / 3 | 30% | High | colon |
| V70 | Male | 45 | TSA | 40% / 3 | 70% | High | colon |
| V72 | Female | 55 | T | 50% / 3 | 80% | Low | rectum |
| V73 | Female | 54 | T | 75% / 3 | 50% | Low | colon |
| V76 | Male | 60 | T | 30% / 3 | 40% | Low | rectum |
| V12 | Male | 53 | T | 60% / 2 | 30% | High | colon |
| V15 | Male | 65 | T | 10% / 1 | 40% | Low | colon |
| V16 | Female | 69 | T | 20% / 3 | 60% | Low | colon |
| V25 | Male | 57 | T | 20% / 2 | 60% | Low | colon |

*T = Tubular, TSA = Traditional serrated; **% of positive tumor cells / staining intensity (0-3); ***Negative

**Supplementary Table S3. Patient characteristics (n=11) and association with immunohistochemical expression of TROP2 and Ki67 in pT1 stage.**

| ID sample | Gender | Age (years) | Histology | TROP2 Score* | Ki67 Score | Localization |
|---|---|---|---|---|---|---|
| 100030_89 | female | 73 | pT1 | 40% / 3 | 30% | Colon sigma |
| 100030_144 | male | 70 | pT1 | 5% /2 | 70% | Colon sigma |
| 100030_263 | female | 53 | pT1 | 70% /3 | 40% | Colon right-side |
| 361/13 1-6 | male | 77 | pT1 | 80% /3 | 60% | Colon right-side |
| 7430/13 1-19 | male | 80 | pT1 | 40% /3 | 80% | Colon sigma |
| 23219/13 1-8 | male | 70 | pT1 | 10% /1 | 30% | Colon right-side |
| 23512/13 1-12 | male | 73 | pT1 | 60% /3 | 40% | Colon ascendent |
| 33516/13 1-10 | male | 75 | pT1 | 5% /1 | 80% | Colon ascendent |
| 4887/16 1-7 | female | 56 | pT1 | 30% /1 | 65% | Colon sigma |
| 100030_943 | female | 68 | pT1 | 35% /2 | 60% | Colon right-side |
| 100030_1038 | male | 68 | pT1 | 5% /1 | 50% | Colon right-side |

*% of positive tumor cells / staining intensity (0-3)

**Supplementary Table S4. Average of reads count obtained for each tissue pair (n=16)**

| Heatmap ID | Cohort | Type of tissue | Total paired - end reads |
|---|---|---|---|
| R01N | CZ | Normal | 46,620,315 |
| R01A | CZ | Adenoma | 47,393,507 |
| R02N | CZ | Normal | 60,338,100 |
| R02A | CZ | Adenoma | 67,024,157 |
| R03N | CZ | Normal | 47,517,187 |
| R03A | CZ | Adenoma | 501,84,951 |
| R04N | CZ | Normal | 47,575,291 |
| R04A | CZ | Adenoma | 51,706,429 |
| R05N | CZ | Normal | 57,623,632 |
| R05A | CZ | Adenoma | 44,545,668 |
| R06N | CZ | Normal | 51,760,500 |
| R06A | CZ | Adenoma | 43,443,514 |
| R07N | CZ | Normal | 45,034,904 |
| R07A | CZ | Adenoma | 50,173,569 |
| R08N | CZ | Normal | 60,101,120 |
| R08A | CZ | Adenoma | 42,515,327 |
| R09A | IT | Adenoma | 44,039,093 |
| R09N | IT | Normal | 64,282,225 |
| R10A | IT | Adenoma | 48,780,993 |
| R10N | IT | Normal | 58,937,614 |
| R11A | IT | Adenoma | 62,027,125 |
| R11N | IT | Normal | 59,177,838 |
| R12A | IT | Adenoma | 57,506,032 |
| R12N | IT | Normal | 55,402,629 |
| R13A | IT | Adenoma | 50,867,452 |
| R13N | IT | Normal | 54,663,873 |
| R14A | IT | Adenoma | 49,867,304 |
| R14N | IT | Normal | 45,320,921 |
| R15A | IT | Adenoma | 47,332,700 |
| R15N | IT | Normal | 59,237,197 |
| R16A | IT | Adenoma | 53,930,082 |
| R16N | IT | Normal | 44,005,481 |

**Supplementary Table S5. Primer pairs (Metabion) of target genes used in this study.**

| Gene ID | Sequence |
|---|---|
| ***MMP1*** | Forward: AGAGCAGATGTGGACCATGC<br>Reverse: TTGTCCCGATGATCTCCCCT |
| ***MMP7*** | Forward: AGTGGTCACCTACAGGATCG<br>Reverse: GGGATCTCTTTGCCCCACAT |
| ***CLDN1*** | Forward: CCCAGTCAATGCCAGGTACG<br>Reverse: CAAAGTAGGGCACCTCCCAG |
| ***CLDN2*** | Forward: GACTCCACTGAGGAACTGCC<br>Reverse: CACAGCTCCCACACTGTCAT |
| ***ETV4*** | Forward: AGTGCCCTACACCTTCAGCAG<br>Reverse: CAGGAACAAACTGCTCATCACTG |
| ***TACSTD2*** | Forward: TCCCCTTTCGGTCCAACAAC<br>Reverse: AAACGATCCCGGGTTGTCAT |
| ***β2-MB*** | Forward: AAGATGAGTATGCCTGCCGT<br>Reverse: CTGCTTACATGTCTCGATCCCA |

**Supplementary Table S6. Primer sequences designed to align *TACSTD2* promoter in 21 CpG sites.**

| | |
|---|---|
| **Pyro Mark CpG Assay** | |
| Forward primer | internal information of the manufacturer Qiagen (Germany) |
| Reverse primer | internal information of the manufacturer Qiagen (Germany) |
| Sequencing primer | internal information of the manufacturer Qiagen (Germany) |
| Sequence to analyze | ACGCGCCAGGTCTGTAGCAGGAGGCCGCGCCGAGGGCGG |
| **CpG Assay 1** | |
| Forward primer | TAAAGAAGAGAGGGAGTGAGAGAA |
| Reverse primer | ACCTCCTACTACAAACCT |
| Sequencing primer | GGAAAGAAAGAAAAGGGA |
| Sequence to analyze | GTCGCGTATAGAGGAGAGCGCGACAGT |
| **CpG Assay 2** | |
| Forward primer | GGAGGAGGAGGGAGTAGTTTT |
| Reverse primer | CCCCACTAATATTTAAATAACACATCC |
| Sequencing primer | GGAGTAGTTTTTTTGTTTTGA |
| Sequence to analyze | CGCGGGCGGCGCAGGGCCGGCTTGGCCTTCCGTGGGACGGGGAGGGGGGC |
| **CpG Assay 3** | |
| Forward primer | GGGGGGAGGGATGTGTTATTTAA |
| Reverse primer | CTCCCTCCCACTCTTATACTCTAC |
| Sequencing primer | GTGTTATTTAAATATTAGTGGGGA |
| Sequence to analyze | CGGTCGGTGGTGGAACCAGCCGGGCAGGTCGGGT |

**Supplementary Table S9. Biological functions of candidate genes.**

| Code | Name | Function |
|---|---|---|
| ***MMP1*** | Matrix metallopeptidase 1 | *MMP1* is a member of family of matrix metalloproteinases (MMPs). Its physiological function is a degrade of extracellular matrix during the development processes, tissue remodeling or metastasis. For enzyme activation is $Ca^{2+}$ reacquired. *MMP1* overexpression is associated with CRC [7, 8]. |
| ***MMP7*** | Matrix metalloproteinase 7 | *MMP7* is a member of family of matrix metalloproteinases (MMPs). It is a proteolytic enzyme, lacking the C-terminal hemopexin domain compared with other family members, it reduces cell adhesion by digestion of extracellular matrix [9] *MMP7* has been studied in the context of CRC [10-12]. |
| ***CLDN1*** | Claudin-1 | *CLDN1* is a transmembrane protein, responsible for formation of tight junctions and contributing to the tight permeability barrier. It is regulated by phosphorylation to increase the tight junction and is found in various tissues, including the intestines, skin, or kidneys [13]. It indirectly affects water homeostasis, ion movements, and seal the gaps between adjacent cells [14]. Dysregulation of *CLDN1* expression has been described in association with several types of cancer, including CRC and chronic inflammatory disease [13]. Indeed, changes in *CLDN1* levels result in weakened intercellular junctions and contribute to increased cell motility, one of the hallmarks of tumor progression [15]. |
| ***CLDN2*** | Claudin-2 | *CLDN2* is transmembrane protein, responsible for cell junction. It increases the permeability; its role is as a pore-forming claudin. Increased permeability promotes the controlled movement of molecules across cell layers [16]. *CLDN2* is tissue-specific, especially in organs where increased permeability is required such as the intestine and the proximal renal tubules [17, 18]. Overexpression of *CLDN2* has been detected in inflammatory bowel disease and cancer and is associated with tumor progression [19]. |
| ***ETV4*** | ETS Variant 4 | *ETV4* is a transcriptional activator of targeted genes involved in cell proliferation (*CCND1, CDK4, CDK6, CCNE1, CUL)* [20] and invasion (*MMP13*) [21], its increased expression was observed in tumor growth, but also in metastatic stages. Due to its involvement in enhancing the invasive properties of cancer cells and malignancies, *ETV4* has already been studied as a potential therapeutic target [21]. In cancer processes, the relationship of downregulation of E-cadherin by *ETV4* was described in the framework of epithelial-mesenchymal transition (EMT). Consequently, this leads to a decrease in cell adhesion characteristics and a shift towards a more mesenchymal and migratory phenotype [22]. |
| ***TACSTD2*** | Tumor-Associated Calcium Signal Transducer 2 also known as TROP2 (Trophoblast Cell-Surface Antigen 2) | Transmembrane glycoprotein, fulfills numerous crucial biological functions, one of which is promoting robust cell-to-cell adhesion [23]. Participates in the differentiation of epithelial cells and the regulation of stem cells due to its surface expression. Its fundamental role is in the calcium signaling [24]. TROP2 is often detected at heightened levels in various cancer types, notably in epithelial-origin carcinomas like CRC. Furthermore, it is linked to tumor advancement and metastasis due to its involvement in intracellular adhesion. Therefore, it is a potential biomarker and therapeutic target in cancer treatment [25, 26]. |
