## Supplementary figures and images for "*TACSTD2* expression marks the early transition to colon adenomas"

### Supplementary Figures

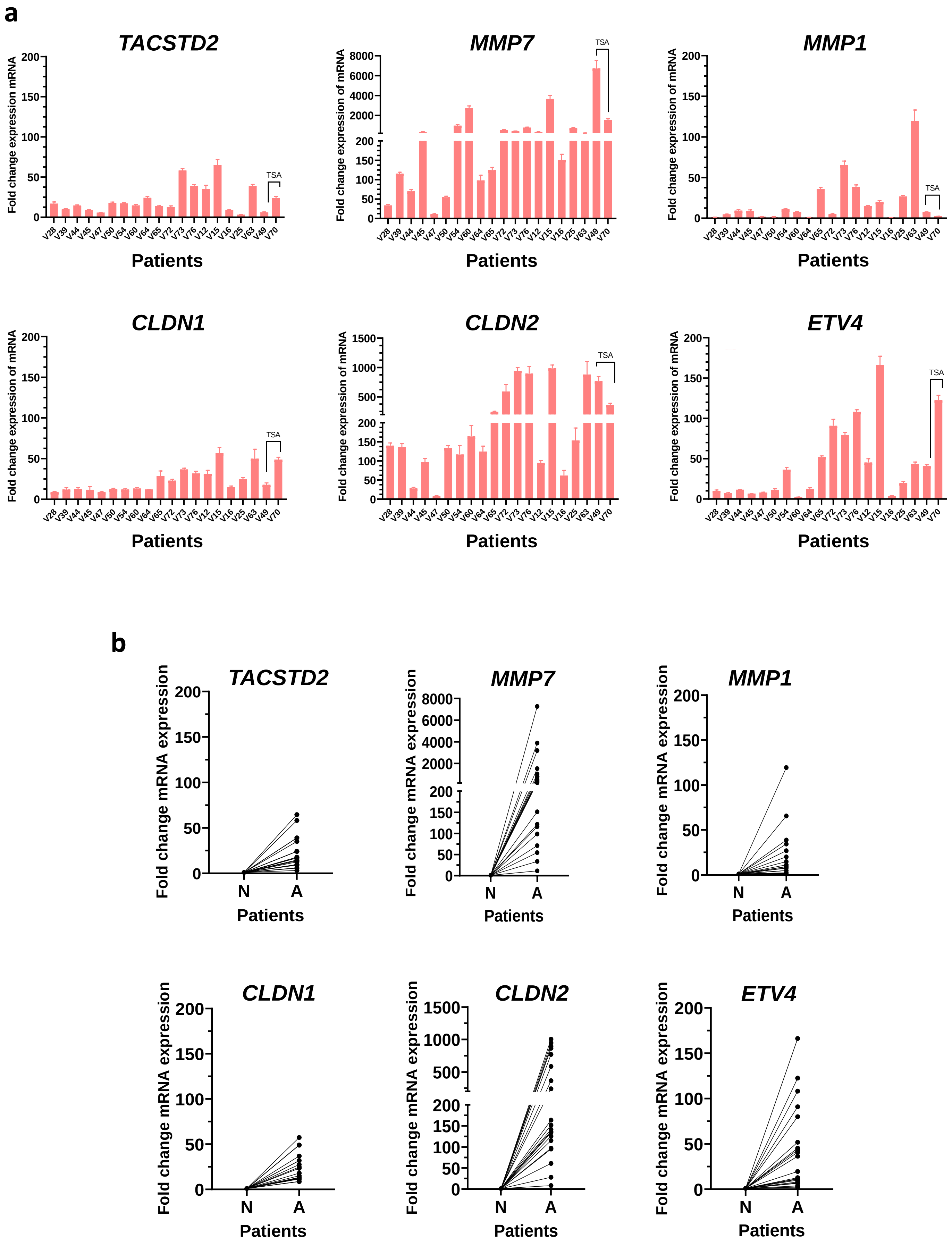


**Supplementary Fig. 1**

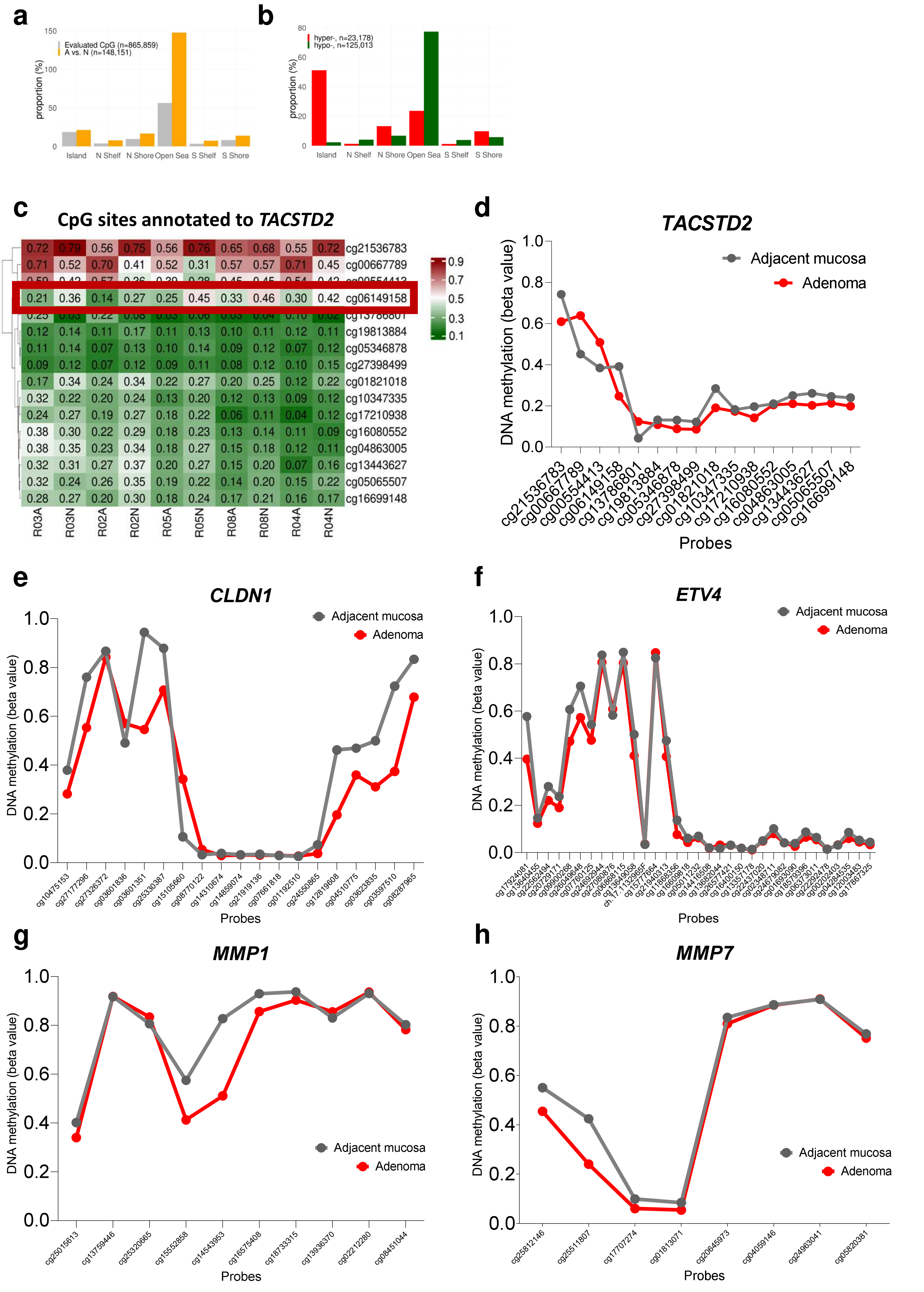


**Supplementary Fig. 2**

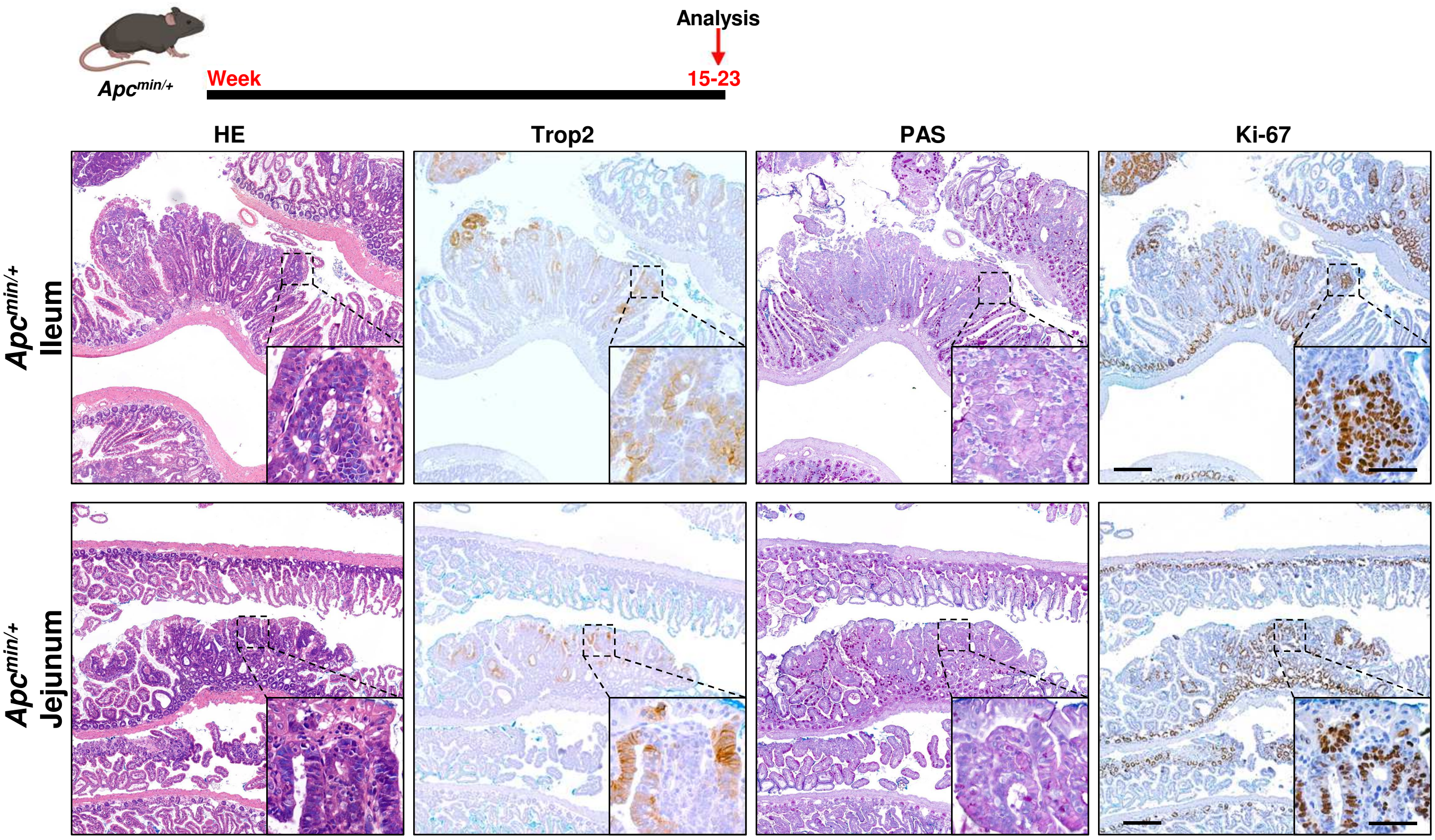


**Supplementary Fig. 3**
